# Deep Reinforcement Learning Enables Better Bias Control in Benchmark for Virtual Screening

**DOI:** 10.1101/2023.11.03.565594

**Authors:** Tao Shen, Shan Li, Xiang Simon Wang, Dongmei Wang, Song Wu, Jie Xia, Liangren Zhang

## Abstract

Virtual screening (VS) has been incorporated into the paradigm of modern drug discovery. This field is now undergoing a new wave of revolution driven by artificial intelligence and more specifically, machine learning (ML). In terms of those out-of-the-box datasets for model training or benchmarking, their data volume and applicability domain are limited. They are suffering from the biases constantly reported in the ML application. To address these issues, we present a novel benchmark named MUBD^syn^. The utilization of synthetic decoys (i.e., presumed inactives) is the main feature of MUBD^syn^, where deep reinforcement learning was leveraged for bias control during decoy generation. Then, we carried out extensive validations on this new benchmark. First, we confirmed that MUBD^syn^ was superior to the classical benchmarks in control of domain bias, artificial enrichment bias and analogue bias. Moreover, we found that the assessment of ML models based on MUBD^syn^ was less biased as revealed by the analysis of asymmetric validation embedding bias. In addition, MUBD^syn^ showed better setting of benchmarking challenge for deep learning models compared with NRLiSt- BDB. Overall, we have proven that MUBD^syn^ is the close-to-ideal benchmark for VS. The computational tool is publicly available for the easy extension of MUBD^syn^.

## 1. Introduction

Virtual screening (VS) is born to simulate the wet-lab high-throughput screening campaigns at significantly low expense of time and money^1^. The target-based method is one of the most promising strategies as it has been validated by a large number of prospective applications^2^. In contrast to the rapid development of new techniques and models, there is great deficiency in the target-specific data committed to training and assessment. For one thing, the data size and diversity of active compounds are extremely limited when novel targets are concerned, resulting in the restricted applicability domain (AD) of ligand-based methods^3^. For another, adequate data for inactive compounds is essential for the simulation of real-world screening scenario. However, the target-specific inactives that have been experimentally validated remain scarce as most of the screening campaigns do not make all negative data accessible^4^. This situation promotes the employment of decoys, i.e., presumed inactives, in the construction of benchmarking datasets for the simulation.

Since Bissantz *et al*. retrospectively compared the performance of different docking approaches on a primitive database in 2000^5^, numerous benchmarking datasets have been published with their construction methodologies extensively studied^3^. For instance, Directory of Useful Decoys (DUD), followed by DUD-Enhanced (DUD-E) has become the golden standard for molecular docking assessment^6,7^. Benchmarking biases including artificial enrichment bias, false negative bias and analogue bias were mitigated by the chemical library-based screening strategies in structure-based virtual screening (SBVS). Solid works regarding bespoke benchmarking datasets for ligand-based virtual screening (LBVS) were also reported, where Maximum Unbiased Validation (MUV) datasets pioneered this field^8^. MUV focuses on optimal embedding of actives in the descriptor-defined chemical space where decoys are uniformly spread. Consequently, two-dimensional (2D) topological bias which causes overestimation of LBVS could be significantly reduced. Inspired by the spatial statistics used in MUV, a series of target-specific benchmarking datasets named Maximal Unbiased Benchmarking Datasets (MUBD)^9^ have been published in the last decade. They are applicable to both LBVS and SBVS methods. Recently, the computational tool named MUBD-DecoyMaker 2.0 was released with a Python-based graphical user interface, which is publicly accessible by the scientific community^10^. By carefully dealing with the inductive bias in the strategy for decoy production^11^, the construction of benchmarking datasets for classical VS methods becomes not that challenging. However, the simple extrapolation of ADs of these classical datasets to the recent VS techniques represented by machine learning (ML) will inevitably introduce new biases^12^. Wallach *et al*. demonstrated that the data clumping in both active set and inactive set causes huge redundancy in the random split of benchmarking datasets for training and validation, thus proposing the asymmetric validation embedding (AVE) for bias detection^13^. Sieg *et al*. discovered that ML approaches are prone to the superficial features of molecular data^14^. This situation was also noted by Chen *et al*.^15^, who named it as “decoy bias”. It is attributed to the obvious distinctions in topological structures between actives and decoys. As a result, the discrimination can be an easy task for deep neural networks. In summary, these benchmarking biases can make ML approaches overestimated in performance assessment but poor in generalization on unseen data.

The decoys of most VS benchmarking datasets are retrieved from chemical libraries including ZINC^16^, ChEMBL^17^ and PubChem^18^. Despite the rapid growth in their data size, these real libraries can never cover entire chemical space where drug-like molecules alone are estimated to exceed 10^6019^, thus making the chemical library-based decoy production strategy hard to make decoys that ideally meet the predefined criteria. The situation is even worse for MUV or the recently published LIT-PCBA^20^ that rigorously requires real compounds with determined bioactivity for a specific target. In fact, this fundamental issue has been recognized by the community. Wallach *et al*., as the pioneers, computationally generated virtual decoys which constituted a DUD- strategy based benchmarking dataset called Virtual Decoys Set (VDS)^21,22^. The virtual decoys synthesized from chemical building blocks show better physicochemical matching and remain topologically dissimilar to ligands. In recent years, the exploitation of synthetic data gains popularity with the emerging of deep generative models^23^. Their applications have rapidly expanded from computer vision to biomedicine, particularly *de novo* drug design^24^, which is in turn leveraged by the VS benchmarking community. Imrie *et al*. innovatively introduced graph-based variational autoencoder to the *in silico* generation of property-matched decoys^25^. While DeepCoy is capable of generating large-scale decoys with less artificial enrichment bias, its model architecture determines that the quality of virtual decoys highly depends on the pairs of molecules for model training. Fortunately, such an issue was circumvented by the progress of constrained molecular generation. In a very recent study, Zhang *et al*. was able to expand the chemical space of target-specific decoy molecules based on a conditional recurrent neural network^26^. With the feature named conformation decoys being integrated, they revealed that the ML-based scoring functions trained on the datasets built by TocoDecoy truly learned implicit physics of protein-ligand interactions. To the best of our knowledge, the previous studies have not made the most of virtual decoys to make ideal benchmarking datasets. Albeit the remarkable progress of methodologies in building generative models for decoys production, the criteria to make unbiased decoys remain the same as the standard of DUD-E, which are physicochemically similar but topologically dissimilar to the ligands. Due to the significant progress towards the multi-objective optimization of molecular properties^27,28^, it is now feasible to generate decoys with additional molecular features. Herein, we utilized deep reinforcement learning (RL) to generate maximal unbiased datasets for VS benchmarking and/or model training. First and foremost, the multiparameter objective (MPO) scoring function of deep generative model named REINVENT^29^ was highly customized to incorporate all the debiasing algorithms from MUBD. Then, the candidate decoys generated by REINVENT were further curated to balance the MPO score and structural diversity. After the development of this new computational tool, we built a series of datasets named MUBD^syn^ and detected their benchmarking biases. We also conducted comparative study to evaluate their feasibility as a benchmark to assess various VS methods such as similarity search, molecular docking and ML-based ones. Particularly, both classical ML models and emerging deep learning models were included, with AVE biases computed to detect the data clumping.

## 2. Material and methods

### 2.1. Data collection and curation

The datasets used as the source or for comparison in this work included Unbiased Ligand Sets/Unbiased Decoy Sets (ULS/UDS)^30^, MUV^8^, DUD-E^7^, DeepCoy^25^, TocoDecoy^26^ and NRLiSt BDB^31^.

In the internal validation, 17 ligand sets from ULS were taken as the input, based on which MUBD^syn^ was made by the pipeline shown in Figure 1A. For comparison, MUBD^real^ was constructed by MUBD-DecoyMaker 2.0^10^ that employs “real” chemical library-based decoy production strategy.

**Figure 1.**
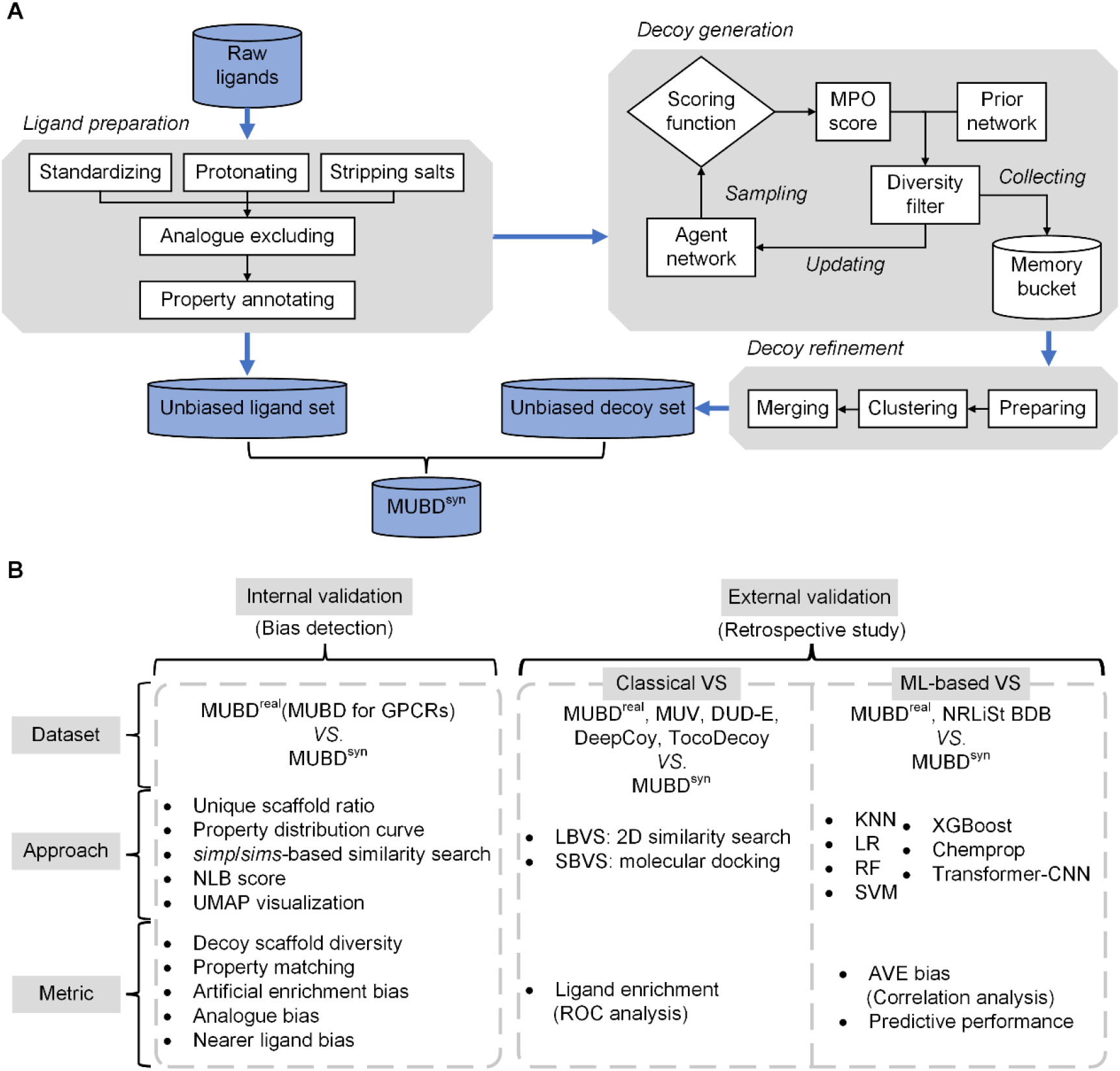
The construction and validation of MUBD^syn^. (A) The pipeline for making MUBD^syn^; (B) The MUBD^syn^ validation consists of “Internal validation” with well-known metrics and “External validation” with classical VS (similarity search and molecular docking) and ML-based VS.

Five cases, i.e., AID 652/Human immunodeficiency virus type 1 reverse transcriptase (HIVRT), AID 712/Heat shock protein HSP 90-alpha (HSP90A), AID 713/Estrogen receptor alpha (ESR1), AID 733/Estrogen receptor beta (ESR2) and AID 810/Focal adhesion kinase 1 (FAK1), were selected for external validation on classical VS methods (i.e., similarity search for LBVS and molecular docking for SBVS). These cases were covered by both MUV and DUD-E, and thus were available for a fair comparative study with MUBD^syn^. The corresponding benchmarking datasets for these cases were then obtained from MUV and DUD-E. MUBD^real^ and MUBD^syn^ for these cases were constructed, with five ligand sets of MUV as input. Additionally, DeepCoy and TocoDecoy for the aforementioned five cases were taken into comparison as well. The benchmarking datasets of DeepCoy were accessed from the published resources without any processing^32^. The benchmarking datasets of TocoDecoy were made with the scripts provided at the GitHub repository^33^ and five ligand sets of DUD-E as input. For TocoDecoy, it should be noted that the grid filter with 300 units was used to refine the candidate decoys, resulting in five TocoDecoy_9W datasets.

In terms of the external validation on ML-based VS methods, ten ligand sets from NRLiSt BDB were taken as the input to make MUBD^syn^ in the way mentioned above, and MUBD^real^ for comparison as well. These ligand sets had sufficient number of diverse ligands after the curation of raw data, and their names were PR_agonist, LXR_alpha_agonist, LXR_beta_agonist, AR_antagonist, PPAR_beta_agonist, PR_antagonist, ER_beta_agonist, ER_alpha_agonist, PPAR_alpha_agonist and PPAR_gamma_agonist.

### 2.2. Ligand preparation

All the ligands were used in the representation of simplified molecular input line entry system^34^ (SMILES). Data curation with MolVS^35^ (version 0.1.1) and Dimorphite-DL^36^ (version 1.3.2) including molecule standardization, salt stripping and protonation at the pH range from 7.3 to 7.5 were performed. To build the unbiased ligand set, “Analogue excluding” and “Property annotating” were performed on the ligand set. The “Analogue excluding” was achieved by the iterative selection of ligands based on the threshold of pairwise similarity between ligands. During the selecting loop, any ligand whose Molecular ACCess System (MACCS) structural keys-based similarity defined by Tanimoto coefficient (Tc) to the referenced one was beyond 0.75 was removed from the raw ligand sets. For “Property annotating”, each ligand was annotated with its raw and linearly normalized values of six properties which were molecular weight (MW), number of rotatable bonds (RBs), number of hydrogen bond donors (HBDs), number of hydrogen bond acceptors (HBAs), net formal charge (nFC) and LogP. The annotated information also included the maximum and minimum values of these properties and pairwise similarity, which were employed to set the training configuration in the next step. RDKit^37^ (version 2020.09.1.0) was used to compute all the properties.

### 2.3. Construction of MPO score

MPO score constitutes the core of cost function of RL in REINVENT^29^. In our specific task for decoy generation, ten scoring components were designed to transform real chemical library-based MUBD decoy screening algorithms into RL-based decoy debiasing algorithms. The structure of our customized MPO score is shown in Figure S1A.

According to the original workflow for MUBD construction^30^, the scoring components included the preliminary ones and the precise ones. The former part was further divided into two major component collections, i.e., components for property filter and components for topology filter. To be more specific about the design of components for property filter, in order to make a single score calculated by each scoring component within the interval of [0, 1], value transformation based on step functions was imposed on the raw scores. Consequently, a generated molecule would be scored 1 by each property component if its calculated physiochemical property was within the range of the corresponding property filter, otherwise it would be scored 0 (Figure S1B). The components for topology filter were designed in a similar manner. Left and right step functions were respectively employed for value transformations to define the high and low thresholds of this filter. Precise scoring components were designed based on two metrics, i.e., *similarity in properties* (*simp*) calculated by the Eq. (1) and *similarity in structure difference* (*simsdiff*) calculated by the Eq. (2):

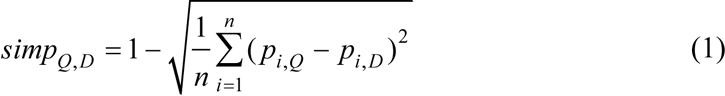

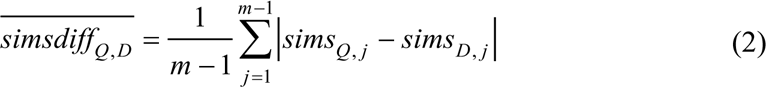

The *simp* measures the similarity in physicochemical properties between the query ligand and the generated decoy. In the Eq. (1), *p* represents the normalized value of physicochemical property, *Q* or *D* is respectively denoted as query ligand or generated decoy, *i* refers to the index of individual property and *n* is the total number of properties. The value of *simp* was directly taken as the output of corresponding scoring component without extra transformation. For *simsdiff*, it was formulated to measure the relative topological difference between query ligand and generated decoy, where the anchor point was defined by the rest of ligands collectively. In the Eq. (2), *similarity in structure* (*sims*) is the Tc of MACCS structural keys-based similarity between two compounds, *j* refers to the index of each remaining ligand and *m* denotes the total number of unbiased ligands. Reverse sigmoid function was imposed on the raw value of *simsdiff* (Figure S1B). Notably, parameters of this transformation function were adjusted to achieve the balance between the return of a specific value extremely close to 1 when *simsdiff* is 0 and a steep drop in the output when *simsdiff* increases. Weighted sum^29^ was taken to organize all the scoring components.

### 2.4. Training REINVENT

The customized training parameters were configured based on the REINVENT community notebook^38^. To be more specific about the parameters of this block, the prior and agent networks were both initialized by the pretrained model named “random.prior.new” provided at the repository. Moreover, the training stopped when it reached 2,000 epochs. For the block of “Diversity Filter”, “Identical Murcko Scaffold” was set as the type of controlling molecular diversity and its bin size was set to 5. Notably, the minimum total score was set to 0.9, which means that only the generated decoy that could achieve an average score over 0.9 was stored in the memory during the training period. Finally, scoring function was configured based on the description above. The parameters not mentioned here were set as default according to the RL notebook.

### 2.5. Decoy refinement

The potential decoy sets were generated ligand by ligand. Concretely, the generative model was trained individually for each ligand and the corresponding decoy set would be collected in the memory when the training was done. As the ratio of decoys to ligands was set to 39 in our previous study^30^, it was essential to refine the potential decoy set before the selection of the most unbiased decoys for each ligand. Curation was firstly conducted to remove invalid SMILES and duplicates from the datasets, followed by molecular clustering based on structural similarity. The agglomerative clustering implementation of scikit-learn^39^ (version 1.2.2) was employed to obtain 39 clusters and then the top-ranking decoys were retrieved from each cluster. Finally, all the selected decoys were annotated with the properties and merged into the unbiased decoy set.

### 2.6. Validation of MUBD^syn^

As shown in Figure 1B, the internal and external validations constituted the full validation scheme. The internal validations focused on the established metrics that detected the benchmarking biases, and the datasets used here all belonged to the MUBD series. In comparison, the external datasets including MUV, DUD-E, DeepCoy, TocoDecoy and NRLiSt BDB were obtained for external validations, wherein the main theme was retrospective study with both classical and ML-based VS methods.

#### 2.6.1. Unique scaffold ratio

The Bemis-Murcko atomic framework^40^ was extracted from each decoy and the duplicates were removed to produce a unique scaffold set. The ratio of scaffold to decoy was calculated as a metric of chemical diversity. Accordingly, a decoy set achieves maximum diversity when the ratio is 1.

#### 2.6.2. simp/sims-based similarity search

Similarity search was implemented in the form of leave-one-out cross-validation (LOO CV) where each query ligand was iteratively selected from the diverse ligand set and pairwise similarity was calculated between the query ligand and the remaining compounds. In terms of artificial enrichment bias, *simp* was calculated to constitute the score list and the label that was true or false (i.e., ligand or decoy) was assigned to the corresponding compound in the score list. Eventually, receiver operating characteristic curve (ROC) was plotted by Matplotlib^41^ (version 3.4.3), and area under curves (AUCs) obtained by scikit-learn were statistically analyzed to get mean(AUCs)s and standard deviations. The diagonal line with AUC equal to 0.5 in ROC analysis signifies random discrimination of samples. Accordingly, the dataset was less biased in term of artificial enrichment if the ROC was closer to the diagonal line or AUC was closer to 0.5. As an intuitive supplement to artificial enrichment bias measurement, physiochemical property matching between the ligand set and the decoy set was also validated in the form of property distribution curve plotted by Matplotlib.

Analogue bias was measured in a similar pattern while the similarity search was based on *sims* between the query ligand and each compound from both ligand and decoy subset. Mean(AUCs)s and standard deviations were also calculated to reflect the extent to which LBVS was distorted by non-uniform compound distribution, and the value of 0.5 denoted that the dataset was free of such bias.

#### 2.6.3. Near ligands bias (NLB) score

This metric has been applied to the comparative study of previous MUBD^9^ with DUD-E and DEKOIS. It is formulated in the Eq. (3) and the Eq. (4):

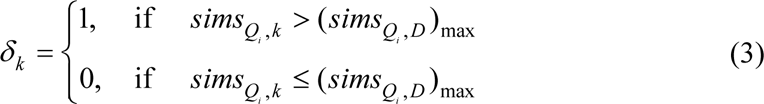

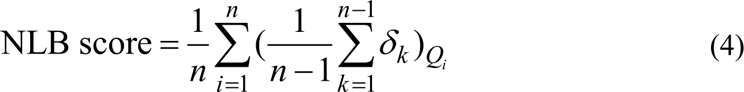

Briefly, let *i* be the index for query ligand *Q* and *δk* be a state scalar for remaining ligand *k*, *δk* is counted 1 when *sims* between query ligand *Qi* and ligand *k* is greater than the maximum *sims* between query ligand *Qi* and each decoy molecule *D*, otherwise 0. Taking *n* as the total number of ligands, formula in brackets computes the percentage of near ligands (NLs) in one iteration of cross validation, and then it is summed and averaged to quantify overall 2D bias.

#### 2.6.4. Uniform manifold approximation and projection (UMAP) visualization

Visualization and analysis of chemical space were realized by UMAP^42^ provided in the umap-learn (version 0.5.3). Both active and decoy molecules were encoded by two kinds of molecular representations. The physicochemical descriptors were six normalized properties including MW, RBs, HBDs, HBAs, nFC and LogP whereas the other representations were MACCS structural keys. Both kinds of representations were embedded into two dimensions and visualized with Matplotlib.

#### 2.6.5. Classical VS methods

LBVS, which adopted the ligand-based similarity search in the form of LOO CV, was performed and ROC analysis was used to evaluate the performance. Two Morgan fingerprints, i.e., ECFP_4 and FCFP_6, were generated for each molecule^43^. Followed by iteratively calculating the topological similarity between query ligand and the remaining compounds, ROC analysis was conducted to get their AUCs. The mean(AUCs) and standard deviation were used to quantify the quality of datasets.

In terms of the SBVS approach, smina^44^, which is a fork of AutoDock Vina^45^, was employed for molecular docking. Five receptor structures including HIVRT (PDB: 3LAN), HS90A (PDB: 1UYG), ESR1 (PDB: 1SJ0), ESR2 (PDB: 2FSZ), FAK1 (PDB: 3BZ3) were directly retrieved from DUD-E and all compounds were prepared with the “Prepare Ligands” module of Discovery Studio^46^ (version 2016) to generate 3D conformers and change ionization states at pH 7.4 prior to molecular docking. With the “autobox” function of smina, each bounding box was automatically defined by the cognate ligand located in the binding site. Parameters not mentioned here were set as default for molecular docking. For ligand enrichment assessment, ROC curves were plotted based on the predicted binding affinity scores and true classes of all the compounds. Moreover, AUCs were also calculated for quantitative comparison.

#### 2.6.6. ML-based VS methods

The MACCS structural keys of each molecule in benchmarking datasets were generated as the input for four classification algorithms including K nearest neighbors (KNN), logistic regression (LR), random forest (RF), and support vector machine (SVM). The training parameters were kept consistent with those provided by Wallach *et al*.^13^, and the models were trained in three-fold cross validation. Two balanced metrics were employed to compare all of the model performance, i.e., MCC and F1 score formulated by Eq. (7) and Eq. (8), respectively:

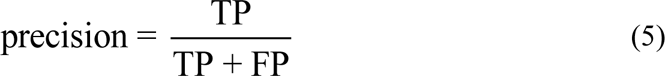

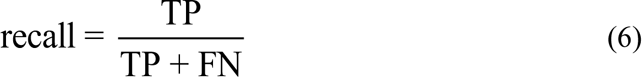

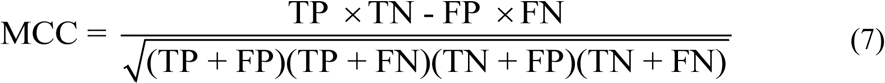

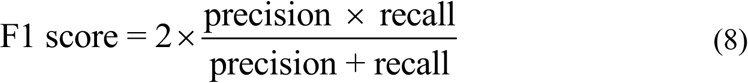

Next, the AVE bias was computed for all datasets. It is comprised of two components that basically leverage the nearest neighbor function of MUV to measure the data clumping:

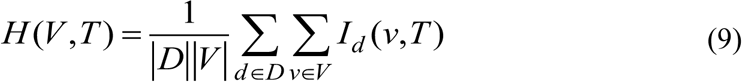

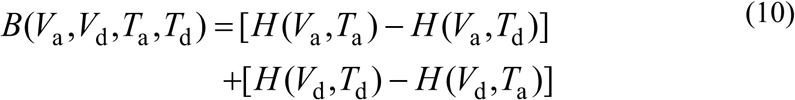

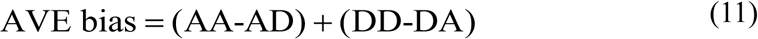

The cumulative nearest neighbor function *H* shown in Eq. (9) describes the extent to which compounds in validation set are similar to those in training set. Eq. (10) further depicts the calculation of AVE bias, and it is simplified in Eq. (11) by the direct replacement of the functional symbols with the pharmacological attributes of corresponding datasets. Herein, “I” and “i” representing inactives in the original equation were replaced with “D” and “d” for decoy molecules in this study. The Pearson correlation coefficient (*ρ*) between MCC and AVE bias was also computed by the SciPy^47^(version 1.7.3).

In the comparative benchmarking, the hyperparameters (“learning_rate”, “max_depth”, “min_child_weight”, “n_estimators” and “subsample”) of eXtreme Gradient Boosting (XGBoost)^48^ on ECFP_4 and the hyperparameters (“depth”, “dropout”, “ffn_hidden_size”, “ffn_num_layers” and “hidden_size”) of Chemprop^49^ were optimized with Hyperopt^50^ (version 0.2.7) via Bayesian Optimization algorithm for 20 iterations, and the objective was set as F1 score given by ten-fold cross-validation on corresponding datasets. The optimized hyperparameters are listed in Table S3. It should be noted that no additional adjustment was carried out to enhance the performance of Chemprop. The default hyperparameters of Transformer-CNN^51^ were used. Three ML models were benchmarked with ten cases from MUBD^syn^ and NRLiSt BDB in the form of five-fold cross-validation.

## 3. Results and discussion

### 3.1. The overview of MUBD^syn^

The pipeline for making MUBD^syn^, briefly shown in Figure 1A, is composed of three submodules. The first module called “Ligand preparation” pre-processes the raw ligands to build the unbiased ligand set, followed by “Decoy generation” module which trains the generative model tuned by RL and collects the potential decoys in the memory, and the last module is called “Decoy refinement” which post-processes potential decoys to build the unbiased decoy set. Compared with the earlier versions of MUBD^10,30^, MUBD^syn^ has three noteworthy features:

1. The unbiased decoy set includes virtual molecules produced by the deep generative model, instead of real compounds screened from chemical libraries;
2. The criteria for an ideal decoy defined in the earlier versions are integrated into a new scoring function for RL to fine-tune the generator;
3. Potential decoys generated by the agent network are further refined to balance the high MPO score and the sufficient chemical diversity.

Accordingly, the aforementioned pitfalls of the chemical library-based decoy selecting strategy were subtly circumvented. For instance, during the first selecting loop of precise filtering in the previous versions, the threshold of *simp*-based physicochemical filtering is lowered in a stepwise manner, i.e., from 0.95 to 0.50 by 0.05, to ensure that each ligand shall have enough decoys for the subsequent *simsdiff*- based topological filtering^30^. Such compromise to the target physicochemical space inevitably brings in artificial enrichment bias. In contrast, MUBD^syn^ is capable of providing enough potential decoys for further filtering. More importantly, the *simp*- based score and the *simsdiff*-based score of a decoy were simultaneously optimized in an unbiased manner. Theoretically, no bias defined by the MUBD rules would be detected in final decoys if such molecules do exist in chemical space. Additionally, the easy expansion of chemical space makes sufficient room for the customized postprocessing of decoys, which may enable the better control of emerging biases including AVE bias^13^, domain bias^14^ and decoy bias^15^.

### 3.2. Internal validation

This kind of validation aims to detect the commonly-observed benchmarking biases. Since the baseline dataset was ULS/UDS, and most metrics here used for comparison originated from the design concepts of MUBD, this section is so-called the internal validation (Figure 1B). The virtual decoy sets of MUBD^syn^ were generated through feeding REINVENT with the corresponding ligand sets, i.e., the ULS dataset in this section.

#### 3.2.1. Chemical scaffold diversity of decoys

It is noted that the average unique scaffold ratio of decoy sets from MUBD^real^ was 0.44 (Figure 2A and Table S1A), which means a unique Bemis-Murcko atomic framework^40^ was shared by more than two decoys. In comparison, there were significant increases in the unique scaffold ratios across all the cases from MUBD^syn^. The average value rose to 0.91 (Table S1B), denoting that almost every molecule in the decoy set had its own unique scaffold. As discussed by Sieg *et al*., chemical diversity deficiency in chemical data leads to the domain bias^14^, which makes the estimated performance of ML models overoptimistic whereas their true power of generalization is in fact not good. It should be noted that the previous work adopted the similar scaffold analysis but only ensured the sufficient chemical diversity of ligand sets in MUBD^9^ whereas neglected it in decoy production. Herein, we have made sure that both scaffolds in the ligand sets and decoys sets from MUBD^syn^ are diverse enough thus the domain bias is well controlled.

**Figure 2.**
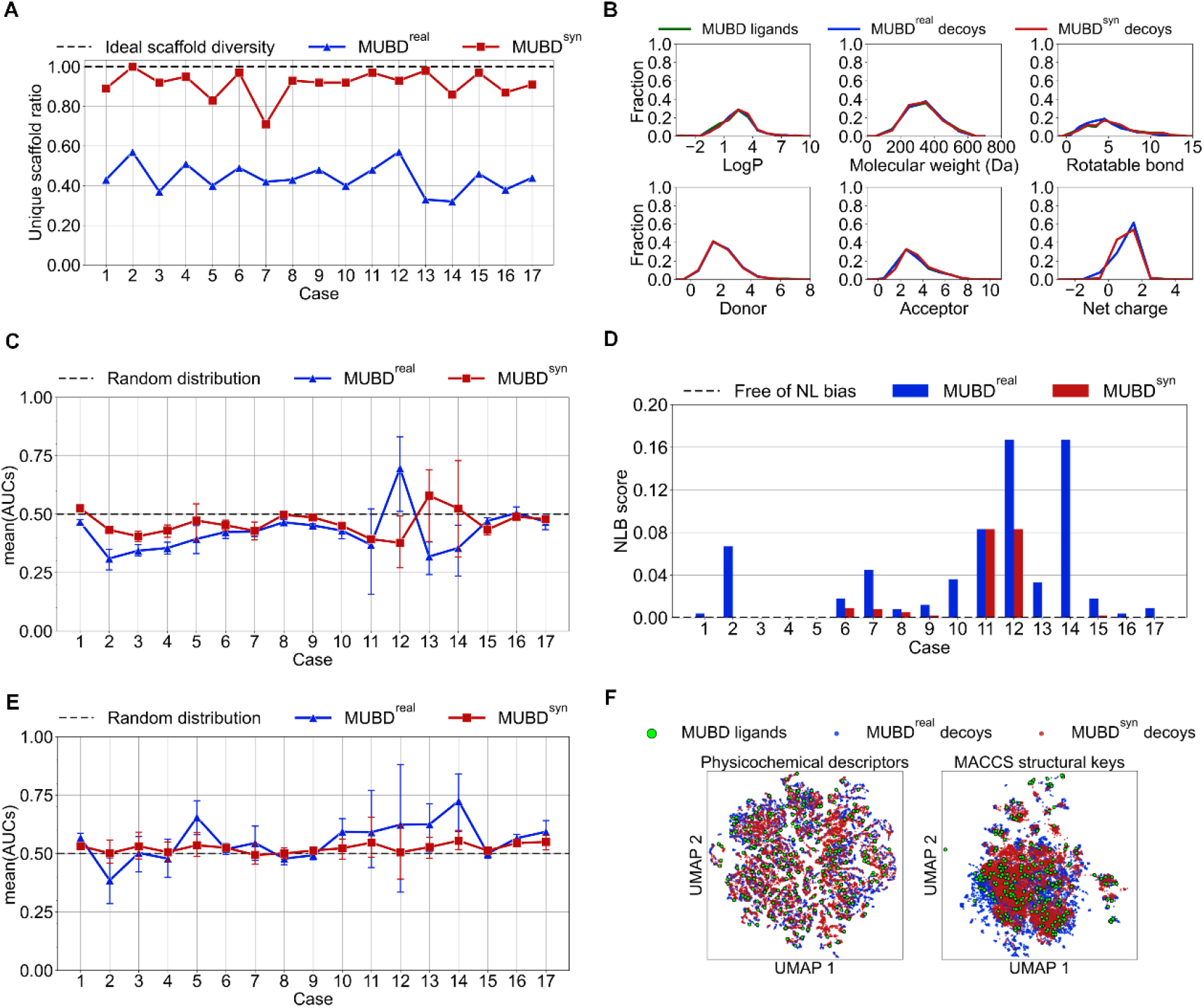
Internal validation for MUBD^syn^ over 17 cases and comparison with MUBD^real^. (A) The unique scaffold ratio of decoys (see Table S1 for case information); (B) The property distribution curves; (C) Performance of the *simp*-based similarity search; (D) NLB score; (E) Performance of the MACCS structural keys *sims*-based similarity search; (F) UMAP visualization of the whole MUBD^syn^ and MUBD^real^ with regard to the chemical space described by physicochemical descriptors (left panel) and MACCS structural keys (right panel), respectively.

#### 3.2.2. Property matching and artificial enrichment bias

As shown in Figure 2B, both decoys from MUBD^real^ and MUBD^syn^ closely matched the unbiased ligands in terms of six physicochemical properties. The excellent property matching was also reflected by the competitive performances of the *simp*- based similarity search for both MUBD^real^ and MUBD^syn^, indicating that artificial enrichment bias was strictly controlled in both MUBDs (Figure 2C). However, it is obvious to find that the mean(AUCs) curve of MUBD^syn^ was closer to the random distribution line. Indeed, the mean(AUCs) of MUBD^syn^ achieved the average value of 0.463, with a minimum of 0.377 and a maximum of 0.579, while that for MUBD^real^ ranged from 0.310 to 0.697 and only achieved the average value of 0.426 (Table S1). This implies that decoys from MUBD^syn^ were more similar to the unbiased ligands in terms of six physicochemical properties, and the rationale behind this improved performance is RL-based generative model continuously generated decoys with high scores.

#### 3.2.3. NLs and analogue bias

NLB score is defined as the average value of the percentage of nearer ligands (NL%) in chemical space, a metric to signify the degree of 2D bias in the benchmarking datasets^9^. Apparently, the overall NLBScore of MUBD^syn^ was much less than that of MUBD^real^ (Figure 2D). It is impressive that 11 out of 17 datasets in MUBD^syn^ were completely free of NL bias whereas only 3 datasets in MUBD^real^ achieved such perfect performance. Accordingly, it is anticipated from these results that the benchmarking performance of LBVS especially for 2D similarity search is less artificially overestimated with MUBD^syn^ than that with MUBD^real^. Consistently, Figure 2E shows that it was more challenging for the similarity search with MACCS structural keys to discriminate the ligands from the decoys of MUBD^syn^ than those of MUBD^real^. That the mean(AUCs) curve of MUBD^syn^ was very close to the random distribution line implies this dataset was almost free of analogue bias. To demonstrate the effects that NLs may exert on 2D similarity search, we analyzed the dataset of PE2R3-AGO. By comparison, we identified the NLB score for PE2R3-AGO from MUBD^real^ was 0.167 while that from MUBD^syn^ was zero (Table S1). Consistently, the 2D similarity search performed much better with MUBD^real^ than MUBD^syn^ for this case, with mean(AUCs) of 0.72 versus 0.55. This case implies that MUBD^real^ is more LBVS favorable and the evaluation outcome based on MUBD^real^ may not be as fair as that based on MUBD^syn^.

#### 3.2.4. Visualization of chemical space

UMAP was utilized to obtain the holistic understanding of interconnections between ligands and decoys (Figure 2F). In terms of the chemical space characterized by physicochemical descriptors, both decoys of MUBD^real^ and MUBD^syn^ uniformly distributed around the ligands, agreeing with their property distribution curves and results of *simp*-based similarity search. Notably, the decoys of MUBD^syn^ were more densely embedded in the adjacent areas of ligands from the point of MACCS structural keys. Individual visualization of two cases from MUBD^real^ (Figure S3B), i.e., SSR2- ANTA and PE2R3-AGO that showed the highest NLBscore (0.167), indeed confirmed that their ligands were nearer to other ligands than the decoys that were sparsely distributed. In contrast, those unbiased but uncharted areas for ideal decoys were occupied by MUBD^syn^, again demonstrating its less NL bias from a brand-new perspective.

#### 3.2.5. Topological features

Looking at the chemical structures of MTR1B-ANTA (Figure 3A), we found that MUBD ^syn^ decoys were bigger than MUBD^real^ decoys in molecular size but more similar to the ligand, thus accounting for the more challenging performance in the *sims*-based similarity search of MUBD^syn^ [mean(AUCs)=0.55] than MUBD^real^ [mean(AUCs)=0.59]. It should be noted that even the most similar decoy of MUBD^syn^ remains below the maximum *sims* threshold of 0.75, satisfying the rule predefined for MUBD decoy.

**Figure 3.**
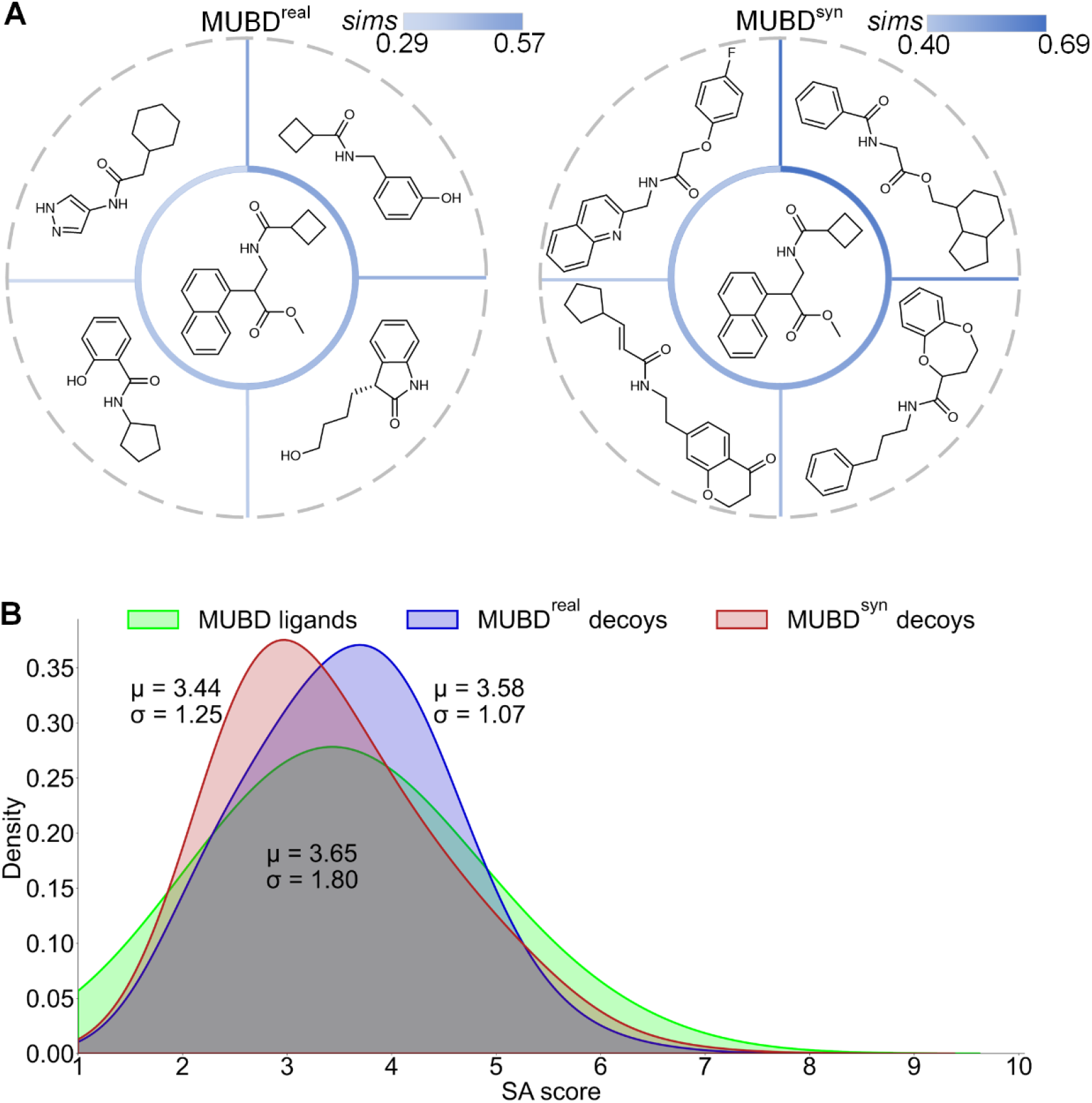
Topological features of MUBD^syn^ decoys and comparison with MUBD^real^ decoys. (A) A glimpse of chemical structures for the case of MTR1B-ANTA. A representative ligand was selected and presented in the central circle. The corresponding MUBD^real^ decoys (left) or MUBD^syn^ decoys (right) were sorted in a descending order by their *sims* values to the ligand, and decoys ranking at the 1^st^, 10^th^, 30^th^, 39^th^ places were selected and presented in the outer ring adjacent to the ligand; (B) The kernel density estimation for SA scores over ligands and the corresponding decoys of MUBD^real^ and MUBD^syn^. These curves were clipped within the range of SA score, i.e., [1,10].

Facilitated by kernel density estimation, we visualized the distribution of topological features that are quantitatively characterized by synthetic accessibility (SA) score^52^ (Figure 3B). The original research pointed out that most of the synthetically accessible molecules fall within the SA score range from two to five. We found that most of the MUBD molecules were synthetically accessible whereas the distribution of MUBD decoys was much sharper and with lower variance. Intriguingly, there was a subtle shift towards lower SA score for the curve of MUBD^syn^ decoys, with the mean value 0.14 less than that of MUBD^real^ decoys, indicating that MUBD^syn^ decoys were less complex in topology.

### 3.3. External validation with classical VS methods

Unlike the internal validation, the external validation was conducted in the form of retrospective study to evaluate the performance of MUBD^syn^ in training and/or benchmarking several kinds of VS methods (Figure 1B). In this section, two classical VS methods including 2D similarity search and molecular docking were applied to five targets including HIVRT, HSP90A, ESR1, ESR2 and FAK1 as case studies for external validation (Figure 4 and Figure S4). The datasets used here can be categorized into two major subsets, i.e., classical datasets of which the data are derived from the real libraries includes MUBD^real^, MUV and DUD-E, and synthetic datasets of which the decoys are made by generative models include MUBD^syn^, DeepCoy and TocoDecoy. It is worth noting that the decoy sets of both DeepCoy and TocoDecoy were made according to the DUD-E ligands. Moreover, the conformation decoys of TocoDecoy were not taken into the comparative study since all the other datasets used in this study only included 2D topological information of the decoys.

**Figure 4.**
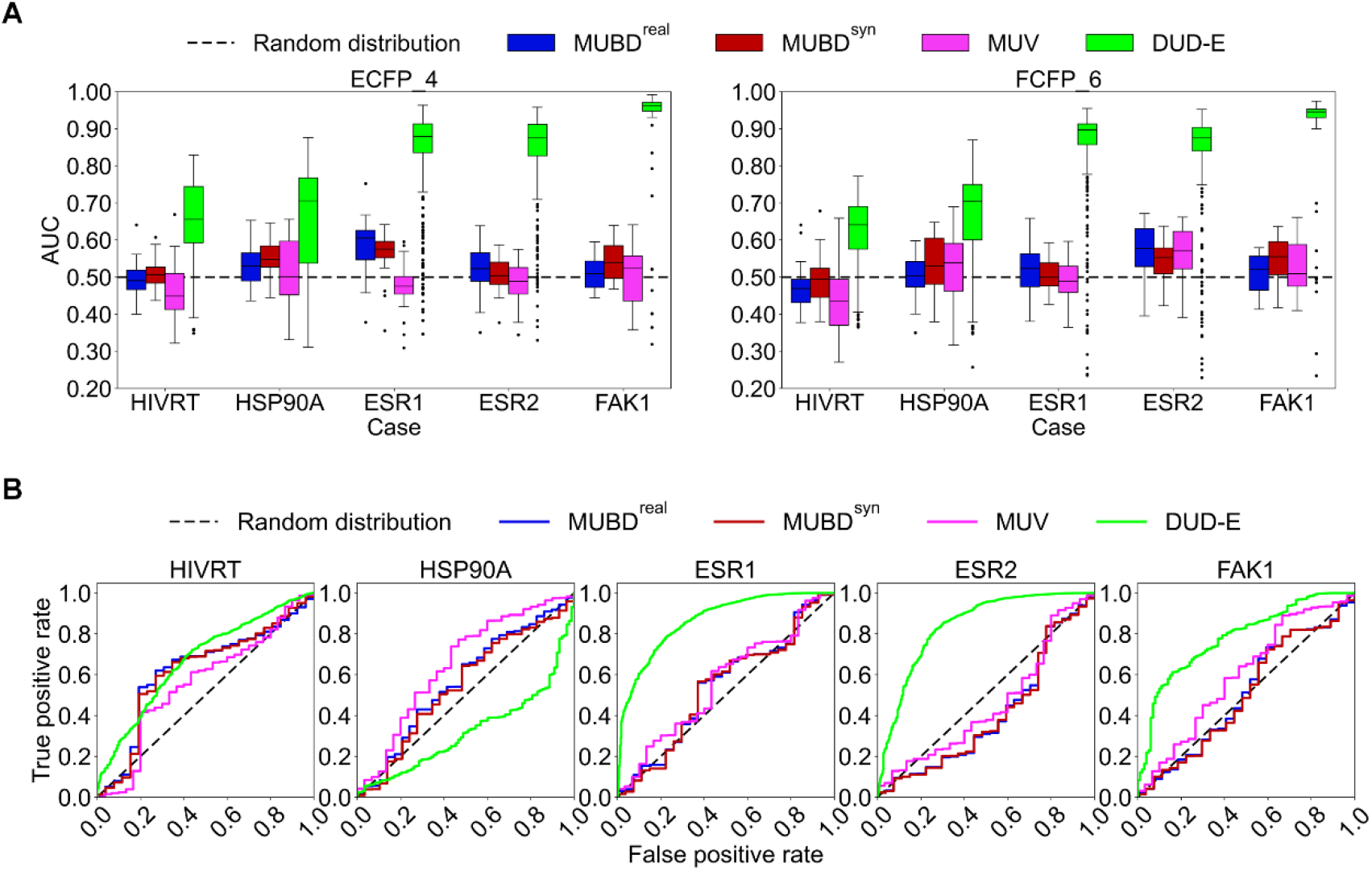
External validation for MUBD^syn^ based on classical VS methods over five cases and comparison with MUBD^real^, MUV and DUD-E. (A) Performance of similarity search (LOO CV) with Morgan fingerprints (ECFP_4 and FCFP_6). The median is represented by the horizontal line of the box; (B) Performance of molecular docking with smina. ROC AUC for each case was computed (Table 1).

**Table 1.** Ligand enrichment measured by ROC AUC with both LBVS approach and SBVS approach over 5 cases in the external validation.

| Case | Approach<sup>a</sup> | Sources of benchmarking datasets |  |  |  |
| --- | --- | --- | --- | --- | --- |
|  |  | MUBD<sup>real</sup> | MUBD<sup>syn</sup> | MUV | DUD-E |
| HIVRT | ECFP_4 | 0.50±0.05 | 0.51±0.04 | 0.46±0.08 | 0.66±0.10 |
|  | FCFP_6 | 0.47±0.06 | 0.49±0.07 | 0.45±0.10 | 0.62±0.09 |
|  | smina | 0.62 | 0.62 | 0.56 | 0.67 |
| HSP90A | ECFP_4 | 0.53±0.05 | 0.55±0.05 | 0.51±0.09 | 0.65±0.15 |
|  | FCFP_6 | 0.50±0.06 | 0.54±0.07 | 0.52±0.10 | 0.66±0.13 |
|  | smina | 0.57 | 0.55 | 0.65 | 0.32 |
| ESR1 | ECFP_4 | 0.59±0.08 | 0.56±0.06 | 0.47±0.06 | 0.84±0.12 |
|  | FCFP_6 | 0.52±0.06 | 0.51±0.04 | 0.48±0.06 | 0.85±0.13 |
|  | smina | 0.54 | 0.54 | 0.56 | 0.86 |
| ESR2 | ECFP_4 | 0.52±0.06 | 0.51±0.04 | 0.49±0.05 | 0.84±0.12 |
|  | FCFP_6 | 0.57±0.08 | 0.55±0.05 | 0.56±0.07 | 0.84±0.13 |
|  | smina | 0.41 | 0.40 | 0.46 | 0.84 |
| FAK1 | ECFP_4 | 0.51±0.04 | 0.55±0.05 | 0.50±0.08 | 0.92±0.14 |
|  | FCFP_6 | 0.51±0.05 | 0.55±0.06 | 0.53±0.07 | 0.90±0.15 |
|  | smina | 0.51 | 0.50 | 0.60 | 0.78 |

<sup>a</sup>For either ECFP\_4 or FCFP\_6 based similarity search, means and standard deviations of AUCs were computed, i.e., presented as mean(AUCs)±std.

#### 3.3.1. Benchmarking of classical LBVS: similarity search

The box plot of similarity search that adopted ECFP_4^43^ shows that the medians of AUCs for both MUBD^real^ and MUBD^syn^ over five cases were all close to the random distribution line (Figure 4A). More importantly, MUBD^syn^ showed more condensed distribution of AUCs than MUBD^real,^ and smaller standard deviations of AUCs except for FAK1 (Table 1). In terms of two chemical library-based datasets, i.e., MUV and DUD-E, MUV was similar to MUBDs whereas decoys of DUD-E were more easily discriminated from its ligands. For example, the similarity search achieved the best AUC of 0.92 on FAK1 case of DUD-E. Even the lowest AUC of 0.65 on HSP90A case of DUD-E was much higher than the highest AUC on MUV (AUC of 0.51 on HSP90A case) or MUBDs (AUC of 0.59 on ESR1 case of MUBD^real^ and AUC of 0.56 on ESR1 case of MUBD^syn^). As mentioned by the developers of DUD-E, 2D VS methods would benefit from the topological difference between ligands and DUD-E decoys due to the design concept, thus the fair benchmarking of LBVS seems impossible^7^. On two synthetic datasets, DeepCoy and TocoDecoy, LBVS achieved similar performance to that on DUD-E (Figure S4A). This outcome was expected, as both datasets follow the principle of DUD-E to make property-matched but topology-distinguished decoys. Nevertheless, we noticed that TocoDecoy was more challenging to enrich except for FAK1 (Table S2), and this observation became more obvious when FCFP_6 was adopted. This may be attributed to the less analogue bias in TocoDecoy, since its topology decoy was further refined by the grid filter based on ECFP^26^. In summary, MUBD^syn^ was comparable to MUBD^real^ and more challenging than MUV while superior to DUD-E and DUD-E-like datasets (i.e., DeepCoy and TocoDecoy) in benchmarking LBVS.

#### 3.3.2. Benchmarking of classical SBVS: molecular docking

For MUBD^real^, MUBD^syn^ or MUV, the ROCs over five cases were all close to the random distribution line (Figure 4B), indicating that it was quite challenging for smina^44^, a renowned software for molecular docking, to enrich the ligands of these benchmarking datasets. Except for HSP90A, discrimination between DUD-E ligands and decoys remained the pretty easy task for smina. It is noteworthy that the AUC for HSP90A of DUD-E was as low as 0.324 (Table 1), and this “anti-screening” phenomenon was also reported by Imrie *et al*. who performed the same retrospective analysis on SBVS^25^, where their reported value of AUC for HSP90A is 0.258.

In terms of two deep learning-generated datasets, DeepCoy and TocoDecoy, the benchmarking results were similar to those of DUD-E, though DeepCoy seemed more challenging on ESR1, ESR2 and FAK1 (Figure S4B). It should be pointed out that TocoDecoy is elaborately designed for ML-based scoring functions. The classical SBVS method used here (i.e., molecular docking with smina) may not reveal its real value because the conformation decoys were not considered here. Nevertheless, DUD-E stays as the golden standard in benchmarking classical SBVS while DeepCoy and TocoDecoy expands the chemical space of DUD-E through synthetic decoys. MUBD^syn^ that was designed according to the same principle as MUBD^real^ may become an alternative in this domain.

### 3.4. External validation with ML-based VS methods

MUBD has been confirmed robust in benchmarking classical VS methods^9^. It was also successfully applied to evaluating the descriptor-based ML models^53^. Herein, we sought to detect ML data redundancy in these datasets and evaluated the plausibility of MUBD^syn^ in training and/or benchmarking ML models. This validation covered both classical ML models based on molecular fingerprints and deep learning models with low-level representations such as text or graph. Normally, MUV should have been used as it was optimized for LBVS. However, its active data size was rather limited, with 30 actives for each target. To ensure the adequate amount and diversity of the active data, we chose NRLiSt BDB^31^ for the validation and selected ten cases (see Table S4 for detailed information), from which the actives were used to make the corresponding datasets of MUBD^syn^ as well as MUBD^real^.

#### 3.4.1. AVE bias and performance correlation analysis

As mentioned above, Wallach *et al*.^13^ provided an insight into the effects of data clumping on ML models training and benchmarking. For a specific molecule in either active set or decoy set, if other molecules, which are similar to that molecule in topological structure, all belong to the same set (active/decoy), there will be data points clumping together when the random partition is adopted. As a result, the data pattern will be memorized rather than generalized by ML models. The zero value of AVE bias indicates that there is no such data redundancy. Their study reported that AVE bias was positively correlated to the performance of ML models constructed with most of the commonly used datasets such as MUV, Tox-21 and PCBA benchmark, pointing out that the random partition is not appropriate for these datasets. We set to calculate AVE bias of MUBD datasets and trained four fingerprint-based ML models with the algorithms of KNN, LR, RF and SVM on MUBD^syn^ and also MUBD^real^ for comparison. Matthews correlation coefficient (MCC) was used as the performance metric due to the imbalance of datasets.

As shown in Figure 5, the AVE bias for each case from MUBD^syn^ was significantly lower than that from MUBD^real^. The average AVE bias over ten cases was as low as 0.10 for MUBD^syn^, and was lower than half of that for MUBD^real^, indicating that the introduction of virtual decoys greatly reduced the data clumping in MUBD. Moreover, the heatmap of model performance reveals that it was more challenging for 1-NN and RF to discriminate ligands from decoys of MUBD^syn^ than MUBD^real^. For example, the average MCCs for MUBD^syn^ given by 1-NN and RF were 0.07 and 0.09, respectively, and this can be considered as random classification. Nonetheless, we deem that relatively close benchmarking performances were achieved on MUBD^syn^ and MUBD^real^ while the former may reflect true capacity of classical ML models due to its lower AVE bias. We noticed that there were moderate correlations between the AVE bias and the predictive performance in both MUBD^syn^ and MUBD^real^, and the highest Pearson correlation coefficient was achieved by SVM (*ρ*=0.87) in MUBD^syn^. Similar correlations were also observed from the separated (AA-AD) term (Figure S5). This indicates that the model assessment could still be overoptimistic when the random partition of MUBD datasets is adopted. Therefore, we suggest that the AVE bias and its correlation coefficient with the model performance should be reported when performing benchmarking with the specific dataset. Moreover, we encourage the employment of either scaffold-based partition^49^ or AVE debiasing algorithms^13^ to further reduce the impact of data clumping in the prospective study.

**Figure 5.**
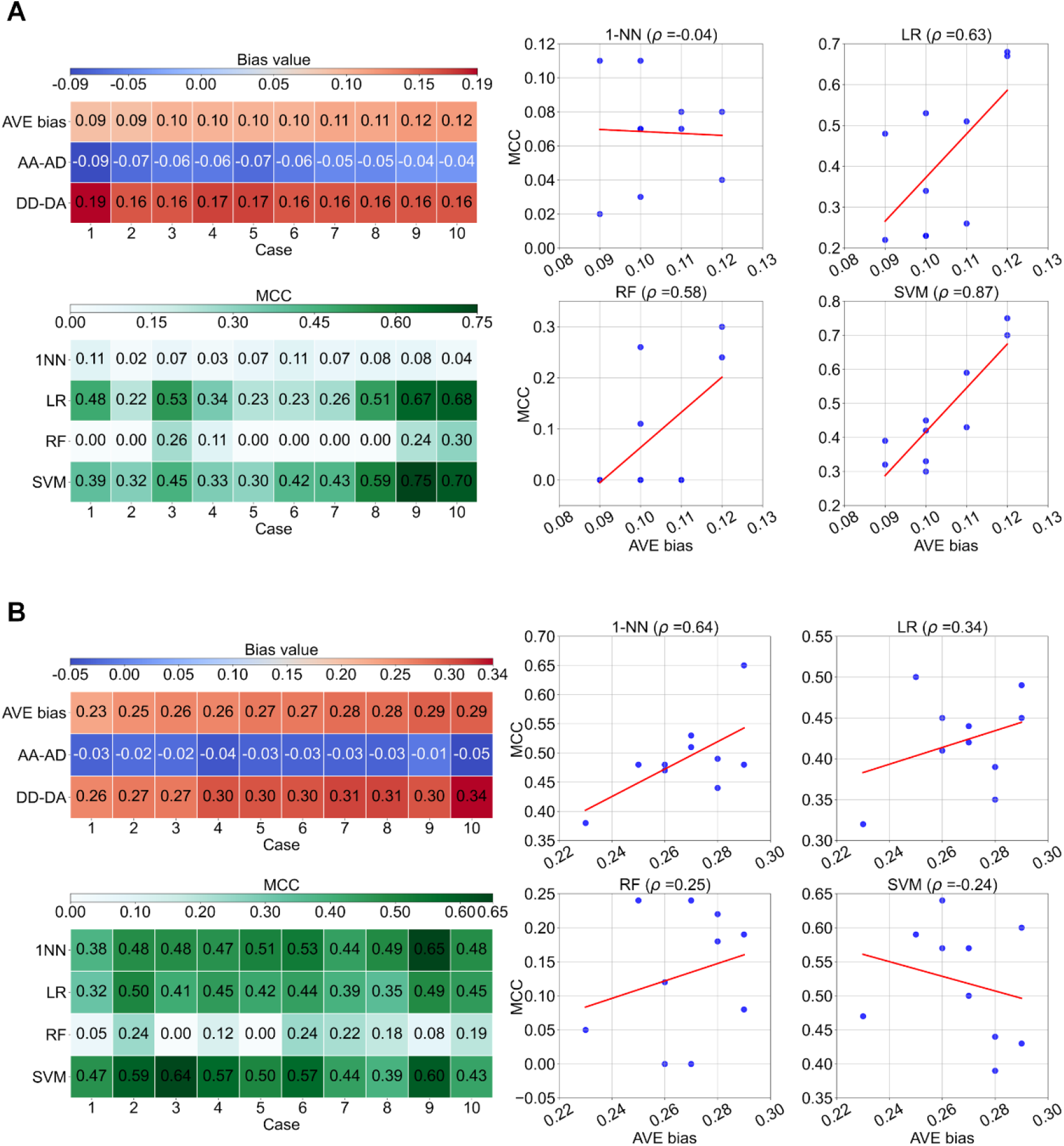
External validation for MUBD^syn^ based on AVE bias and performance correlation analysis. (A) Performances and biases of four ML models trained on MUBD^syn^. Top left: AVE bias and its two decomposed terms for each case. Bottom left: Performances of four ML models measured by MCC (three-fold cross validation). Right: Pearson correlation coefficients (*ρ*) between AVE bias and MCC for each case. The data points were fit with linear regression implemented by regplot function of the seaborn package; (B) Performances and biases of four ML models trained on MUBD^real^ in comparison with MUBD^syn^.

#### 3.4.2. ML-based VS benchmarking

In this section, three representative ML models were assessed with MUBD^syn^ (ten cases from NRLiSt BDB). Specifically, XGBoost on ECFP_4 was set as the baseline representing traditional ML methods while Chemprop and Transformer-CNN, belonging to deep learning models that take 2D molecular graph or SMILES as input, were set as the baseline for advanced ML. It should be noted that the hyperparameters of XGBoost and Chemprop were tuned with Hyperopt based on the same settings (see Table S3 for the optimized hyperparameters) whereas Transformer-CNN was directly used for benchmarking as suggested by its authors^51^. Since there were small datasets in MUBD^syn^ (i.e., fewer than 100 active molecules), we used the form of five-fold cross-validation and average performance to ensure robustness (Figure 6, Figure S6 and Table S4). This form was applied to all the cases to make sure of consistency throughout the whole calculation.

**Figure 6.**
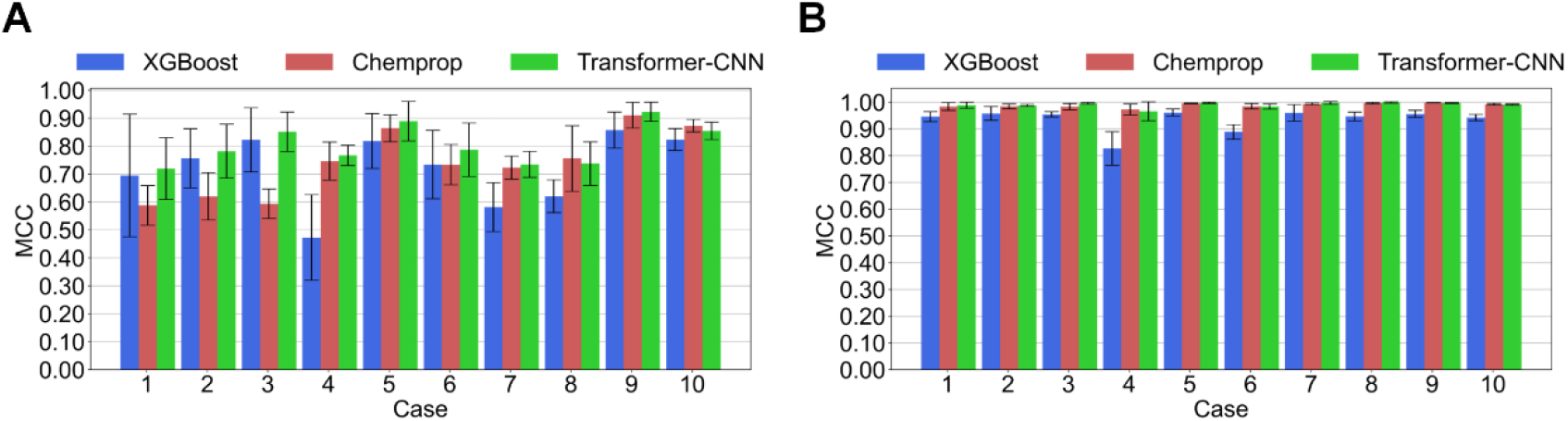
External validation with the benchmark on three representative ML models. Model performance was measured by MCC in the form of five-fold cross-validation. (A) MUBD^syn^; (B) NRLiSt BDB.

For the benchmarking study with MUBD^syn^, advanced ML models (Chemprop and Transformer-CNN) generally outperformed the traditional model (XGBoost) when the size of the active set was larger than 70, while the performances of two deep learning models got very close when the size of active set was larger than 120 (case 8, 9 and 10). It is worth noting that XGBoost was significantly better than Chemprop but worse than Transformer-CNN when evaluated with three cases including PR_ago (case 1), LXR_alpha_ago (case 2) and LXR_beta_ago (case 3). We found that the numbers of actives of these cases were no more than 70. Moreover, the ratio of decoys to actives was kept 39 in MUBD, resulting in data imbalance. As pointed out by the authors of Chemprop^49^, their D-MPNN may underperform traditional ML models if the datasets are small and/or data classes are extremely imbalanced. As such, Chemprop did not perform well in the above mentioned three cases. In comparison, the superiority of Transformer-CNN for small datasets can be attributed to its informative SMILES- embedding derived from a pretraining task about SMILES canonicalization. We also observed the great deviation of performance in the cross-validation on PR_ago (case 1), especially for XGBoost (more than 25% of the average). As expected, the deviation decreased with the growth of data size. For example, the deviations of MCCs on three models were all less than 5% of the average for PPAR_gamma_ago (case 10).

In terms of the benchmark (Figure 6B) NRLiSt BDB, whose decoy sets were made based on the principle of DUD-E, is in fact not a good benchmark for ML models. In all of ten cases, the discrimination of ligands from decoys seemed extremely easy for deep learning models, because even the traditional ML model, i.e., XGBoost achieved excellent classification performance with the average MCC reaching 0.93. With such a benchmarking outcome, advanced ML models poorly distinguished themselves from traditional models. This may impair the interest of the VS community in method development, as it may underestimate the value of new methodology. In fact, this issue has long been acknowledged in other disciplines. For example, the widely used benchmarking dataset named CIFAR^54^ for computer vision has various subsets compiled at different difficulty level to satisfy diverse benchmark demands. As far as this ML-based VS benchmarking was concerned, MUBD^syn^ outperformed traditional benchmark such as NRLiSt BDB when used for evaluating novel ML models due to its appropriate level of benchmark difficulty, and it is encouraging to see that MUBD^syn^ offers a solution to address the aforementioned issue in the field of VS benchmarking.

## 4. Conclusions

VS techniques are constantly advancing, thanks to the evolution of computational theories coupled with the explosive growth in computing capacity^55^, so should be the benchmarking methodologies and databases. In this study, we utilized deep reinforcement learning to make next-generation MUBD. The fundamental update is the revolution of decoy production strategy, which has switched from the chemical library-based screening to the objective-oriented generation. We performed thorough validations on the new datasets named MUBD^syn^. The internal validation has demonstrated its superiority to the previous MUBD in benchmarking bias control. For example, the scaffold diversity of decoy molecules has been significantly improved and NL bias has been almost eliminated in MUBD^syn^. Notably, the UMAP technique has enabled the global visualization of the chemical space where decoys of MUBD^syn^ are embedded more sufficiently and uniformly. The external validation has further confirmed that the application of MUBD^syn^ is not limited to the classical VS methods but extended to emerging ML methods including deep learning. We firstly showed that the AVE bias has been greatly reduced in this new MUBD, thus the ligand enrichment performance of ML models would be less artificially inflated by the data clumping. Furthermore, the comparative benchmark covering both traditional ML model and deep learning models highlighted the importance of compiling the VS benchmarking datasets at an appropriate difficulty level. MUBD^syn^ is challenging enough to present the superiority of advanced ML models while not that challenging to make all models indistinguishable.

The rise of deep learning has boosted the development of computational tools for drug discovery. However, several eye-catching studies^56–58^ that report the successful use of artificial intelligence to discover promising leads all rely on the in-house data (released or not) with limited size for model training. This again emphasizes that the data mining should be the essential work of ML for drug discovery. The constant progress of MUBD is committed to alleviating data deficiency in biomedicine. Compared with DeepCoy and TocoDecoy that have taken advantage of the virtual decoys to make DUD-E-like decoys, we have proved that MUBD^syn^ has broader AD due to its unique debiasing algorithms, and we deem that it will be widely used by the VS community in the future.

This work focuses on the *in silico* augmentation of negative data for VS methods while the shortage of diverse and high-quality positive data remains a concern that should be addressed in the future development of MUBD. In conclusion, our MUBD^syn^ offers a novel way to implement MUBD theory and will be applied to the construction of large-scale VS benchmarking platforms. We posit that MUBD^syn^ will accelerate the computer-aided drug discovery through more robust VS tools.

## ASSOCIATED CONTENT

### Data and Code Availability

The Python and bash scripts, along with the detailed instructions of MUBD^syn^ are available at the GitHub repository (https://github.com/taoshen99/MUBDsyn). In order to ensure reproducibility of all the validations performed in this work, we also provided the guiding notebooks. The template config used for model training will be generated during the implementation of the test case. All the datasets curated and made in this work are available at the Zenodo dataset (https://doi.org/10.5281/zenodo.7943200), which can facilitate the reproduction of all experiments and benchmark of various VS methods.

### Supporting Information

Case study on the optimization of MPO score for MUBD^syn^, supplementary tables and figures of the internal and external validation (Supporting_Information.docx).

## AUTHOR INFORMATION

### Corresponding Authors

*E-mail: J.X., S.W. and D. W. Address: Jie Xia, Song Wu and Dongmei Wang, Institute of Materia Medica, Chinese Academy of Medical Sciences and Peking Union Medical College, Beijing 100050, China

### Conflicts of Interest

The authors declare no conflicts of interest.

### Author Contributions

The project was led by Jie Xia receiving support from Song Wu and Dongmei Wang. The main code was developed by Tao Shen, and necessary modifications were made by Jie Xia and Shan Li. Overall computational experiments were jointly designed by Tao Shen and Jie Xia, and were performed by Tao Shen. The analysis of the results was jointly performed by Tao Shen and Jie Xia. Invaluable feedbacks of the code and experiments were provided by Xiang Simon Wang, Dongmei Wang, Song Wu and Liangren Zhang. The first draft was written by Tao Shen and improved by Jie Xia. The final version of the manuscript was reviewed by all the authors.

### ORCID

Tao Shen: 0009-0005-7086-8803

Jie Xia: 0000-0002-9567-3763

## Acknowledgments

This study was funded by the CAMS Innovation Fund for Medical Sciences (Grant No. 2021-I2M-1-069), the National Science and Technology Major Projects for Major New Drugs Innovation and Development (Grant No. 2018ZX09711001-012-003) and the Program for Foreign Talent of Ministry of Science and Technology of the People’s Republic of China (Grant No. G2021194015L).

## Notes

https://github.com/taoshen99/MUBDsyn

https://doi.org/10.5281/zenodo.7943200

### Competing Interest Statement

The authors have declared no competing interest.

