## Supplementary material for "Deep Reinforcement Learning Enables Better Bias Control in Benchmark for Virtual Screening": Figure S1-6; Table S1-4

Dr. Jie Xia

Institute of Materia Medica

Chinese Academy of Medical Sciences

No. 2 Nanwei Road

Beijing 100050, China

### Table of Contents


S1 Case study on the optimization of MPO score for MUBD$^{syn}$ 3

S2 Internal validation 6

S3 External validation with classical VS methods 10

S4 External validation with ML-based VS methods 12

References 15

#### S1 Case study on the optimization of MPO score for MUBD$^{syn}$

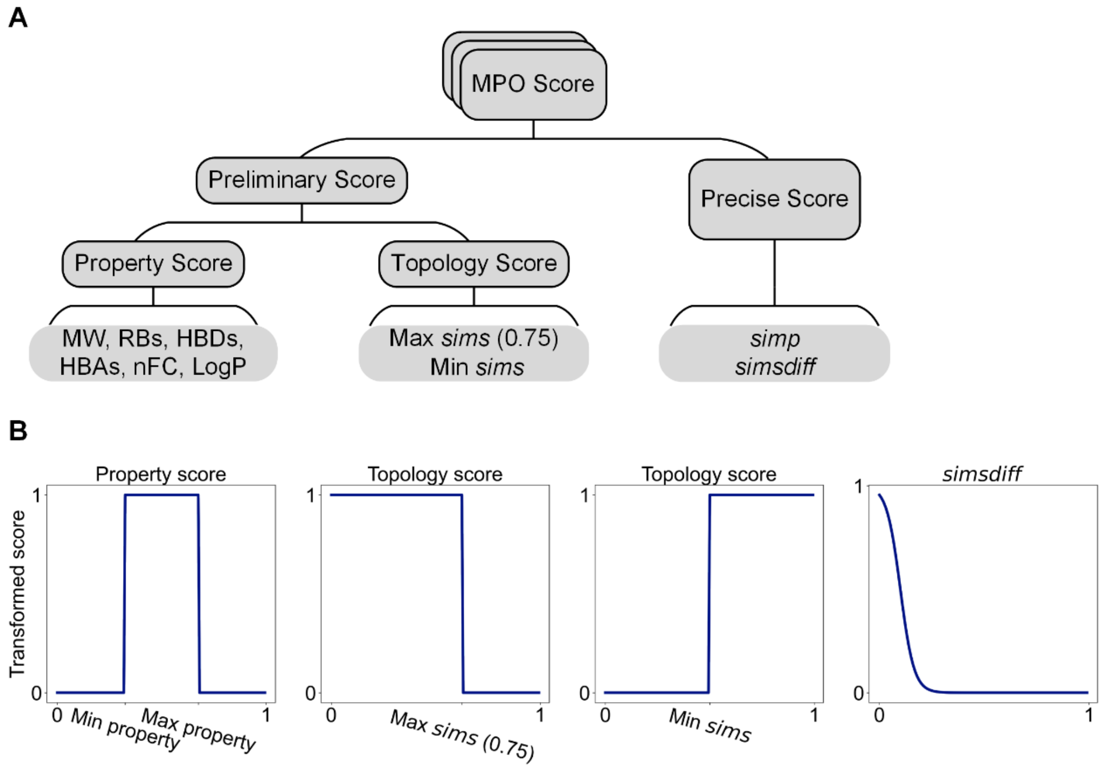


**Figure S1** The construction of MPO score. (A) Ten scoring components of MPO score include MW, RBs, HBDs, HBAs, nFC, LogP, Max *sims*, Min *sims*, *simp* and *simsdiff*. All the components are derived from the filtering algorithms of MUBD-DecoyMaker 2.0 and transformed so as to constitute the scoring function of REINVENT; (B) Transformation functions for scoring components. Left: Step function to transform scaled physicochemical property of decoys to property scoring components. Central left: Left step function to transform max *sims* between decoys and each ligand to topology scoring components. Central right: Right step function to transform min *sims* between decoys and each ligand to topology scoring components. Right: Reverse sigmoid function to transform *simsdiff* score.

It is the high multiparameter objective (MPO) score (Figure S1A) that constantly rewards the agent network to generate more desired molecules, and the training stage of virtual decoy generation for melatonin receptor type 1B antagonists (MTR1B-

ANTA) from Unbiased Ligand Set/Unbiased Decoy Set (ULS/UDS)[1] was taken as a demonstration.

For scoring components based on "preliminary filtering", any undesired decoy will be scored zero if its physicochemical properties or topological similarity to each ligand exceed the preset thresholds. Transformed scores of the first five components including molecular weight (MW), number of rotatable bonds (RBs), number of hydrogen bond donors (HBDs), number of hydrogen bond acceptors (HBAs), net formal charge (nFC), increased rapidly during the initial 400 training steps, indicating that the agent network quickly learned to sample property-matched decoys (Figure S2). At the end of the training stage, all the scoring curves converged to high levels which satisfied the threshold of 0.9 as preset for the memory unit. We noticed that the transformed LogP ended up in a moderate score of 0.73. It is probably the algorithm that adopts the Crippen approach[2] for LogP computation in RDKit made the convergence of LogP more complicated than that of other properties. The training curves of topological filters showed that maximum *similarity in structure* (max *sims*) between the sampled molecule and each ligand was basically within the desired interval while most of the initially sampled molecules did not satisfy the preset minimum *similarity in structure* (min *sims*) of 0.29, which meant that the initial model (trained for less than 400 steps) was inclined to produce molecules which were topologically different from MTR1B-ANTA ligands.

For the last two scoring components constituting "precise filtering", no obvious progress was observed after about 400 steps of training, and transformed scores

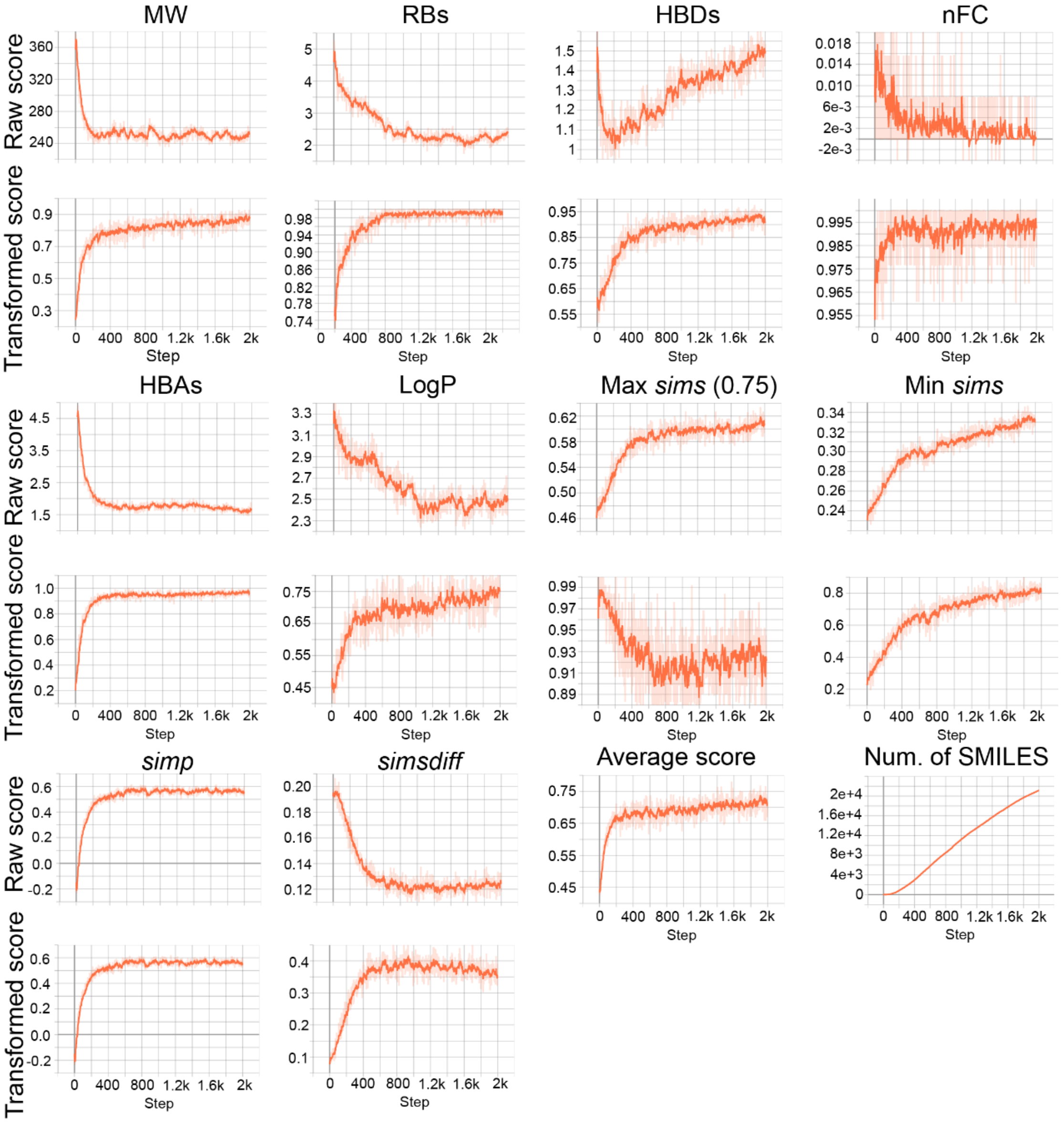


**Figure S2** Optimizing stage of MTR1B-ANTA decoy generation. Tensorboard was used to plot and smooth the training curves of ten scoring components. The average score and accumulated number of candidate SMILES found at each step were also recorded.

eventually converged to the moderate levels, i.e., 0.55 for *similarity in properties* (*simp*) and 0.36 for *similarity in structure difference* (*simsdiff*). The improvement of these scores would be our directions for further development of MUBD$^{syn}$.

In the topic of multi-objective reinforcement learning (RL), the trajectory of optimization is often biased to the easy part of objectives while the complicated part is often less focused, which can be exacerbated by imbalanced weights[3,4]. In principle, autotuning should be performed on the weight of each objective in RL. However, it was computationally expensive to determine the tailored weight distribution for each model, thus each objective was assigned equal weight in this study. Nonetheless, this case showed that the average score (0.71) of the batch sampled at the final step was acceptable and the model collected 21,199 potential decoys which were fairly enough for further refinement.

**S2 Internal validation**

Figure S3 shows the individual chemical space of each case through dimensionality reduction technique implemented by uniform manifold approximation and projection[5] (UMAP). In general, visualizations through case by case agreed with those on the whole datasets. Figure S3A showed the good property matching between ligands and decoys of MUBD. The decoy data points clustered around the ligand data points and formed the different subgroups. This data structure could not be seen in the left panel of Figure 2F. The topological differences of decoys between $MUBD^{real}$ and $MUBD^{syn}$ are shown in Figure S3B. We observed that the red data points representing decoys of $MUBD^{syn}$ were more densely clumped and closer to the ligand data points compared with the blue data points representing decoys of $MUBD^{real}$. As discussed in the main text, such an observation became clearer when the differences in nearer ligands bias (NLB) scores between $MUBD^{real}$ and $MUBD^{syn}$ increased (see cases PER2R3-AGO, SSR2-ANTA and 5HT1F-ANTA that had the greatest differences of NLB score). It indicated that $MUBD^{syn}$ had fewer nearer ligands and its decoys had more shared substructures with ligands, agreeing with the conclusions drawn from the section "3.2.5 Topological features".

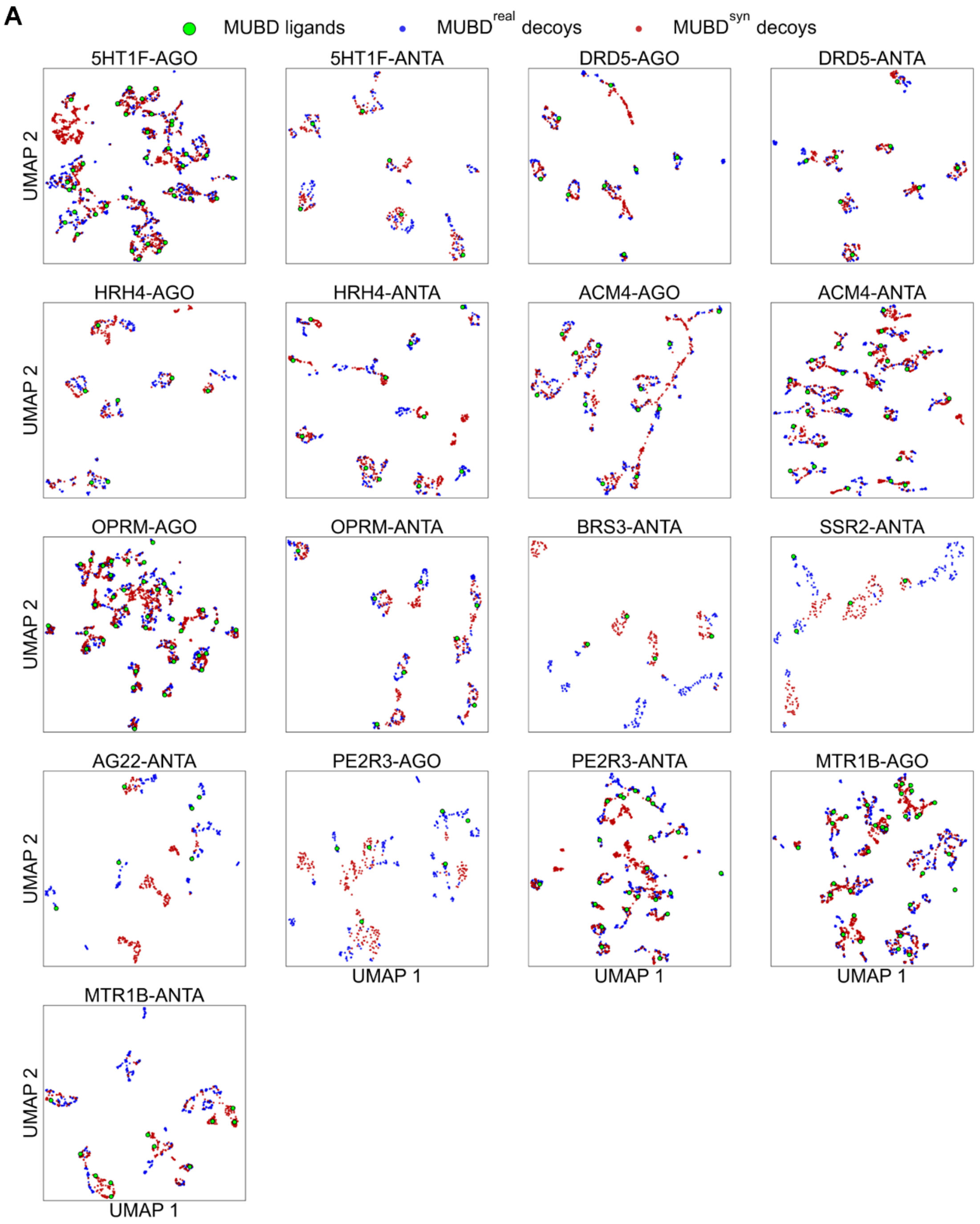

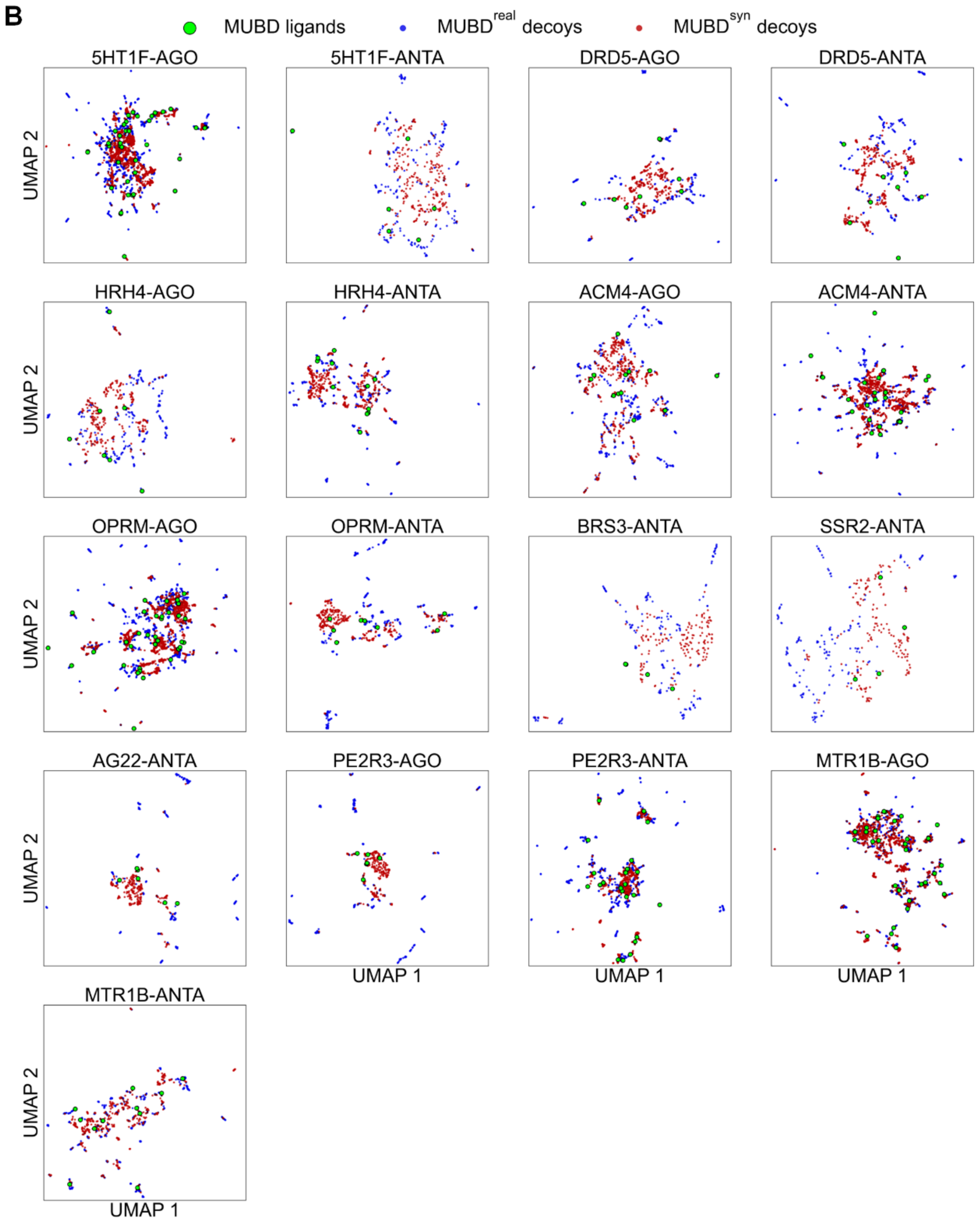


**Figure S3** UMAP visualization of each individual dataset in MUBD$^{real}$ or MUBD$^{syn}$. (A) The chemical space described by physicochemical descriptors; (B) The chemical space described by MACCS structural keys.

**Table S1** Four quantitative metrics used in the internal validation of 17 cases from ULS/UDS. (A) Benchmarking biases in $MUBD^{real}$; (B) Benchmarking biases in $MUBD^{syn}$.

| | | **A** $MUBD^{real}$ | | | | **B** $MUBD^{syn}$ | | | |
|---|---|---|---|---|---|---|---|---|---|
| Index | Case | Unique scaffold ratio of decoys | NLB score | Mean(AUCs)±std (*simp*) | Mean(AUCs)±std (*sims*) | Unique scaffold ratio of decoys | NLB score | Mean(AUCs)±std (*simp*) | Mean(AUCs)±std (*sims*) |
| 1 | 5HT1F-AGO | 0.43 | 0.004 | 0.47±0.03 | 0.57±0.06 | 0.89 | 0.001 | 0.53±0.04 | 0.53±0.05 |
| 2 | 5HT1F-ANTA | 0.57 | 0.067 | 0.31±0.05 | 0.39±0.12 | 1.00 | 0.000 | 0.43±0.01 | 0.50±0.06 |
| 3 | DRD5-AGO | 0.37 | 0.000 | 0.34±0.03 | 0.50±0.11 | 0.92 | 0.000 | 0.41±0.03 | 0.53±0.08 |
| 4 | DRD5-ANTA | 0.51 | 0.000 | 0.35±0.04 | 0.48±0.12 | 0.95 | 0.000 | 0.43±0.04 | 0.51±0.06 |
| 5 | HRH4-AGO | 0.40 | 0.000 | 0.39±0.08 | 0.66±0.10 | 0.83 | 0.000 | 0.47±0.11 | 0.54±0.07 |
| 6 | HRH4-ANTA | 0.49 | 0.018 | 0.42±0.05 | 0.52±0.04 | 0.97 | 0.009 | 0.45±0.04 | 0.52±0.02 |
| 7 | ACM4-AGO | 0.42 | 0.045 | 0.43±0.04 | 0.54±0.14 | 0.71 | 0.008 | 0.43±0.06 | 0.49±0.08 |
| 8 | ACM4-ANTA | 0.43 | 0.008 | 0.47±0.03 | 0.48±0.07 | 0.93 | 0.005 | 0.50±0.03 | 0.50±0.06 |
| 9 | OPRM-AGO | 0.48 | 0.012 | 0.45±0.02 | 0.49±0.06 | 0.92 | 0.002 | 0.49±0.02 | 0.51±0.04 |
| 10 | OPRM-ANTA | 0.40 | 0.036 | 0.43±0.04 | 0.59±0.10 | 0.92 | 0.000 | 0.45±0.02 | 0.52±0.06 |
| 11 | BRS3-ANTA | 0.48 | 0.083 | 0.37±0.19 | 0.59±0.18 | 0.97 | 0.083 | 0.39±0.01 | 0.55±0.10 |
| 12 | SSR2-ANTA | 0.57 | 0.167 | 0.70±0.16 | 0.62±0.30 | 0.93 | 0.083 | 0.38±0.12 | 0.51±0.12 |
| 13 | AG22-ANTA | 0.33 | 0.033 | 0.32±0.08 | 0.62±0.14 | 0.98 | 0.000 | 0.58±0.22 | 0.53±0.06 |
| 14 | PE2R3-AGO | 0.32 | 0.167 | 0.36±0.14 | 0.72±0.16 | 0.86 | 0.000 | 0.52±0.27 | 0.55±0.05 |
| 15 | PE2R3-ANTA | 0.46 | 0.018 | 0.47±0.04 | 0.50±0.05 | 0.97 | 0.002 | 0.43±0.06 | 0.51±0.03 |
| 16 | MTR1B-AGO | 0.38 | 0.004 | 0.50±0.08 | 0.57±0.05 | 0.87 | 0.000 | 0.49±0.03 | 0.54±0.03 |
| 17 | MTR1B-ANTA | 0.44 | 0.009 | 0.47±0.05 | 0.59±0.08 | 0.91 | 0.000 | 0.48±0.02 | 0.55±0.05 |

#### S3 External validation with classical VS methods

**Table S2** Ligand enrichment measured by the area under curve of receiver operating characteristic curve (ROC AUC) of both ligand-based virtual screening (LBVS) and structure-based virtual screening (SBVS) over 5 cases in the external validation.

| Case | Approach[a] | Sources of benchmarking | |
|---|---|---|---|
| | | DeepCoy | TocoDecoy |
| HIVRT | ECFP_4 | 0.61±0.08 | 0.60±0.08 |
| | FCFP_6 | 0.64±0.08 | 0.56±0.08 |
| | smina | 0.60 | 0.57 |
| HSP90A | ECFP_4 | 0.61±0.11 | 0.60±0.09 |
| | FCFP_6 | 0.61±0.09 | 0.57±0.07 |
| | smina | 0.40 | 0.43 |
| ESR1 | ECFP_4 | 0.80±0.14 | 0.79±0.13 |
| | FCFP_6 | 0.84±0.13 | 0.81±0.12 |
| | smina | 0.70 | 0.83 |
| ESR2 | ECFP_4 | 0.80±0.14 | 0.79±0.13 |
| | FCFP_6 | 0.82±0.13 | 0.78±0.12 |
| | smina | 0.69 | 0.81 |
| FAK1 | ECFP_4 | 0.87±0.14 | 0.90±0.12 |
| | FCFP_6 | 0.88±0.16 | 0.88±0.16 |
| | smina | 0.67 | 0.78 |

[a]For either ECFP_4 or FCFP_6 based similarity search, means and standard deviations (std) of AUCs were computed, i.e., presented as mean(AUCs)±std.

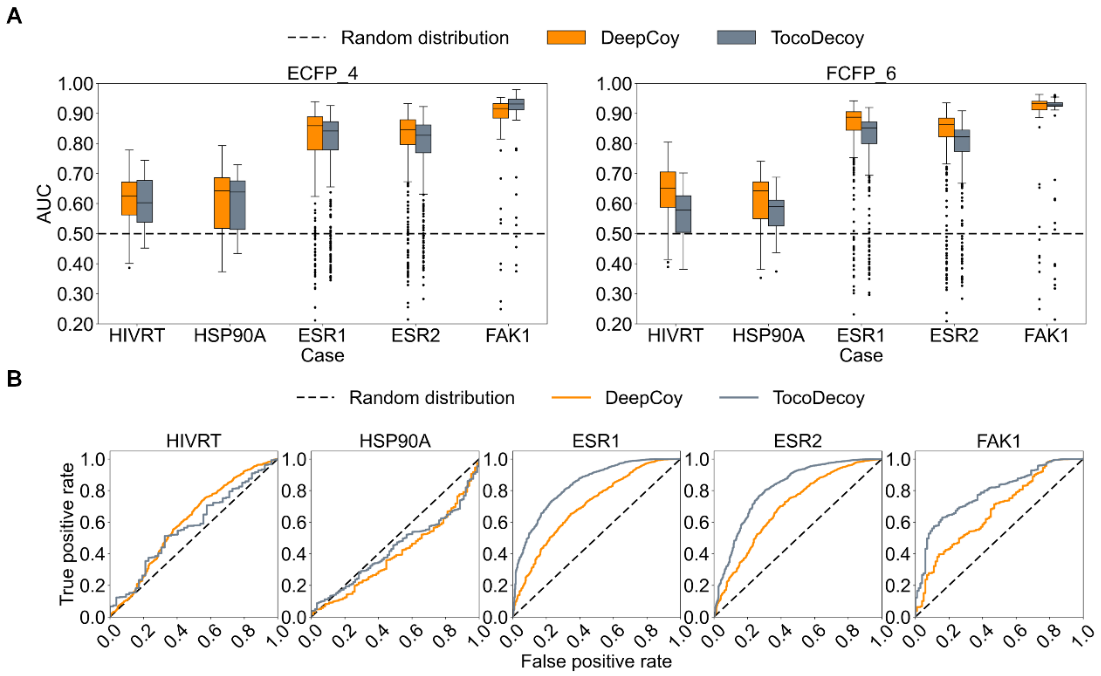


**Figure S4** External validation based on classical VS methods over five cases and comparison with DeepCoy and TocoDecoy. (A) Performance of similarity search with Morgan fingerprints. The median is represented by the horizontal line of the box; (B) Performance of molecular docking with smina. ROC AUC for each case was computed.

#### S4 External validation with ML-based VS methods

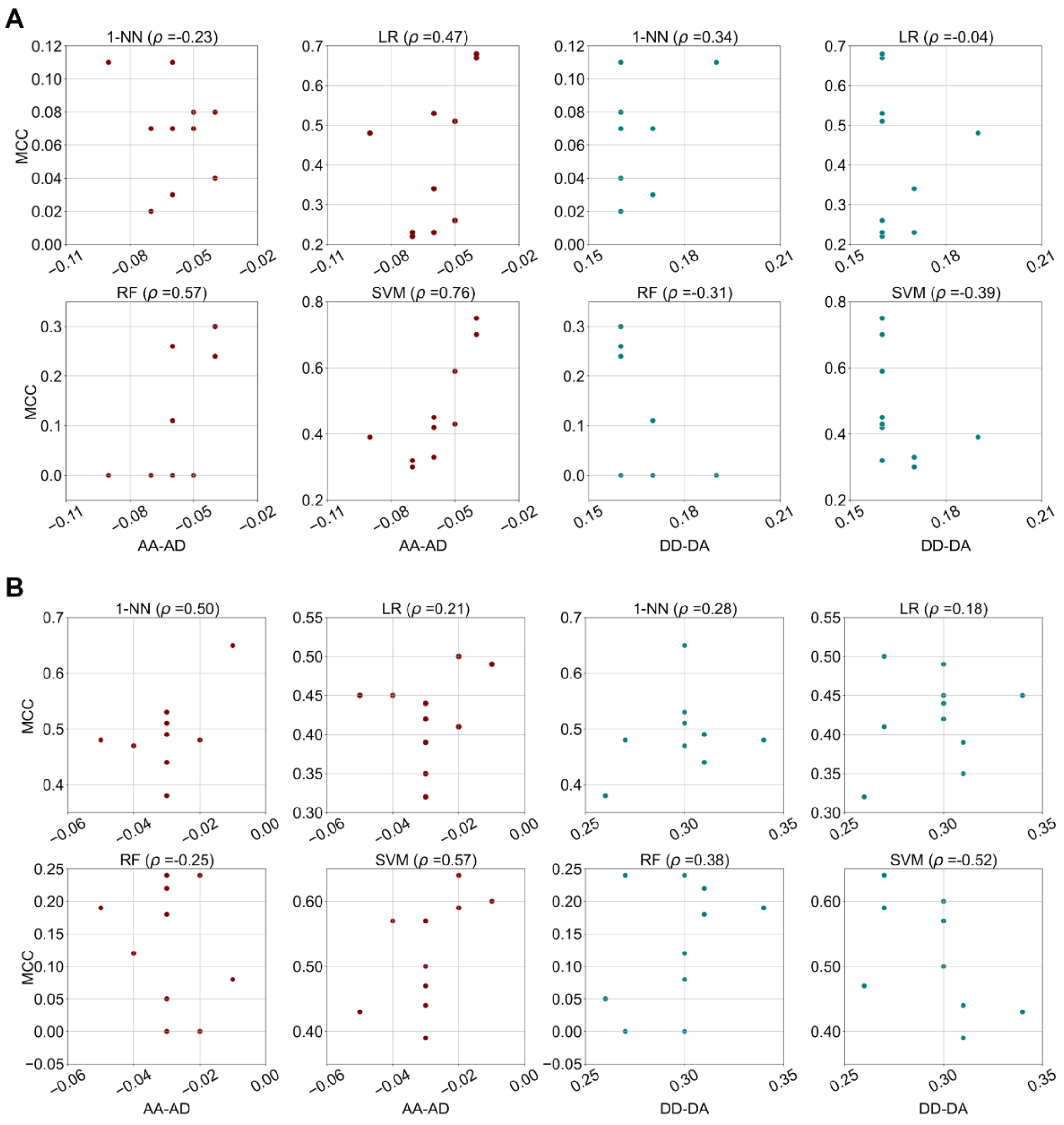


**Figure S5** Pearson correlation coefficients ($\rho$) between separated terms of asymmetric validation embedding (AVE) bias and the predictive performances measured by Matthews correlation coefficient (MCC) of four machine learning (ML) models over ten cases from NRLiSt BDB. (A) Four ML models trained and validated on $MUBD^{syn}$. Left: (AA-AD) term. Right: (DD-DA) term; (B) Four ML models trained and validated on $MUBD^{real}$. Left: (AA-AD) term. Right: (DD-DA) term.

**Table S3** The hyperparameters of XGBoost combined with ECFP_4 fingerprints and Chemprop optimized for the whole dataset of MUBD$^{syn}$ and NRLiSt BDB.

| Model | Dataset | | Model | Dataset | |
|---|---|---|---|---|---|
| XGBoost | MUBD$^{syn}$ | NRLiSt BDB | Chemprop | MUBD$^{syn}$ | NRLiSt BDB |
| learning_rate | 0.05 | 0.05 | depth | 3 | 3 |
| max_depth | 13 | 13 | dropout | 0.4 | 0.05 |
| min_child_weight | 0.85 | 0.85 | ffn_hidden_size | 900 | 1500 |
| n_estimators | 136 | 136 | ffn_num_layers | 1 | 3 |
| subsample | 0.86 | 0.86 | hidden_size | 900 | 1500 |

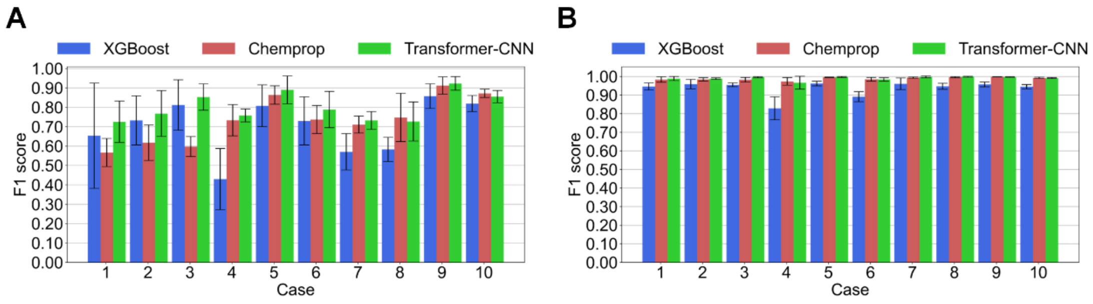


**Figure S6** External validation based on three representative ML models. Model performance was measured by F1 score in the form of five-fold cross-validation. (A) MUBD$^{syn}$; (B) NRLiSt BDB.

**Table S4** Predictive performance measured by F1 score and MCC of XGBoost, Chemprop and Transformer-CNN over ten cases from NRLiSt BDB in the external validation. All the evaluations followed the five-fold cross-validation. The dataset size and AVE bias are also listed for each case. (A) Predictive performance in $MUBD^{syn}$. The cases were sorted by the number of actives in an ascending order. (B) Predictive performance in NRLiSt BDB. The cases are sorted by the case order of $MUBD^{syn}$.

| **A** | | | | $MUBD^{syn}$ | | | | | |
|---|---|---|---|---|---|---|---|---|---|
| | | | | XGBoost | | Chemprop | | Transformer-CNN | |
| Index | Case | Number of actives (decoys) | AVE bias | F1 score[a] | MCC[a] | F1 score[a] | MCC[a] | F1 score[a] | MCC[a] |
| 1 | PR_ago | 47 (1833) | 0.09 | 0.65±0.27 | 0.69±0.22 | 0.57±0.07 | 0.59±0.07 | 0.73±0.11 | 0.72±0.11 |
| 2 | LXRα_ago | 66 (2574) | 0.10 | 0.73±0.13 | 0.76±0.11 | 0.62±0.09 | 0.62±0.08 | 0.77±0.12 | 0.78±0.10 |
| 3 | LXRβ_ago | 70 (2730) | 0.10 | 0.73±0.13 | 0.76±0.11 | 0.62±0.09 | 0.62±0.08 | 0.77±0.12 | 0.78±0.10 |
| 4 | AR_ant | 78 (3042) | 0.11 | 0.43±0.16 | 0.47±0.15 | 0.73±0.08 | 0.75±0.07 | 0.76±0.03 | 0.77±0.04 |
| 5 | PPARβ_ago | 96 (3744) | 0.11 | 0.81±0.11 | 0.82±0.10 | 0.86±0.05 | 0.86±0.05 | 0.89±0.07 | 0.89±0.07 |
| 6 | PR_ant | 103 (4017) | 0.09 | 0.73±0.12 | 0.73±0.12 | 0.74±0.07 | 0.73±0.07 | 0.79±0.09 | 0.79±0.10 |
| 7 | ERβ_ago | 105 (4095) | 0.10 | 0.57±0.09 | 0.58±0.09 | 0.71±0.04 | 0.72±0.04 | 0.73±0.05 | 0.73±0.05 |
| 8 | ERα_ago | 121 (4719) | 0.10 | 0.58±0.06 | 0.62±0.06 | 0.75±0.12 | 0.76±0.12 | 0.73±0.10 | 0.74±0.08 |
| 9 | PPARα_ago | 150 (5850) | 0.12 | 0.86±0.06 | 0.86±0.06 | 0.91±0.04 | 0.91±0.05 | 0.92±0.03 | 0.92±0.03 |
| 10 | PPARγ_ago | 219 (8541) | 0.12 | 0.82±0.04 | 0.82±0.04 | 0.87±0.02 | 0.87±0.02 | 0.86±0.03 | 0.85±0.03 |

| B | | | | NRLiSt BDB | | | | | |
|---|---|---|---|---|---|---|---|---|---|
| | | | | XGBoost | | Chemprop | | Transformer-CNN | |
| Index | Case | Number of actives (decoys) | AVE bias | F1 score[a] | MCC[a] | F1 score[a] | MCC[a] | F1 score[a] | MCC[a] |
| 1 | PR_ago | 269 (12371) | 0.57 | 0.95±0.02 | 0.95±0.02 | 0.98±0.01 | 0.98±0.01 | 0.99±0.01 | 0.99±0.01 |
| 2 | LXRα_ago | 258 (12676) | 0.50 | 0.96±0.03 | 0.96±0.03 | 0.98±0.01 | 0.98±0.01 | 0.99 | 0.99 |
| 3 | LXRβ_ago | 373 (17058) | 0.51 | 0.96±0.01 | 0.95±0.01 | 0.98±0.01 | 0.98±0.01 | 1.00 | 1.00 |
| 4 | AR_ant | 226 (10229) | 0.47 | 0.83±0.06 | 0.83±0.06 | 0.97±0.02 | 0.97±0.02 | 0.97±0.03 | 0.97±0.04 |
| 5 | PPARβ_ago | 906 (38722) | 0.50 | 0.96±0.01 | 0.96±0.01 | 1.00 | 1.00 | 1.00 | 1.00 |
| 6 | PR_ant | 531 (23613) | 0.50 | 0.89±0.03 | 0.89±0.03 | 0.98±0.01 | 0.98±0.01 | 0.98±0.01 | 0.98±0.01 |
| 7 | ERβ_ago | 392 (14209) | 0.64 | 0.96±0.03 | 0.96±0.03 | 0.99 | 0.99 | 1.00±0.01 | 1.00±0.01 |
| 8 | ERα_ago | 434 (16938) | 0.63 | 0.95±0.02 | 0.95±0.02 | 1.00 | 1.00 | 1.00 | 1.00 |
| 9 | PPARα_ago | 1401 (58387) | 0.48 | 0.96±0.01 | 0.96±0.01 | 1.00 | 1.00 | 1.00 | 1.00 |
| 10 | PPARγ_ago | 1820 (43746) | 0.46 | 0.94±0.01 | 0.94±0.01 | 0.99 | 0.99 | 0.99 | 0.99 |

[a]For either F1 score or MCC, means and standard deviations (std) of the metric were computed, i.e., presented as mean±std. Note that the standard deviations less than 0.01 were omitted.
